# AEBP2-Directed H3K27me2 Defines a Specific Vulnerability in EZH2-mutant Lymphoma

**DOI:** 10.1101/2025.10.14.682307

**Authors:** Daniel Angelov, James Nolan, Grainne Holland, Darragh Nimmo, Molly Davies, Craig Monger, Eugene Dillon, Marlena Mucha, Cheng Wang, Dáire Gannon, Eimear Lagan, Andrew Malcolm, Lu Yang, April Chen, Elisabeth Vandenberghe, Andrew Flaus, Chun-Wei Chen, Gerard L. Brien, Eric Conway, Adrian P. Bracken

## Abstract

The catalytic subunit of Polycomb Repressive Complex 2 (PRC2), EZH2, is recurrently mutated in 25% of diffuse large B-cell lymphomas (DLBCL), causing increased H3K27me3 and decreased H3K27me2 levels. EZH2 inhibitors provide clinical benefit, but resistance frequently develops, highlighting the need for alternative therapeutic targets. Here, we identify the PRC2 accessory protein AEBP2 as a specific genetic dependency in EZH2-mutant DLBCL. While AEBP2 acts through PRC2, its essential role is surprisingly independent of canonical H3K27me3-mediated gene silencing. Instead, AEBP2 functions within a PRC2.2 complex lacking JARID2, using its zinc-finger domains to sample intergenic chromatin to sustain H3K27me2. Notably, loss of *AEBP2* or *NSD2* caused contrasting changes in intergenic H3K27me2 levels, driving sensitivity or resistance to PRC2 inhibitors, respectively. Our findings identify AEBP2-PRC2.2-maintained intergenic H3K27me2 as a therapeutic vulnerability in EZH2-mutant DLBCL and highlight dysregulated H3K27me2 as an underappreciated form of PRC2 dysfunction in cancer, with important therapeutic implications.

## INTRODUCTION

Diffuse large B-cell lymphoma (DLBCL) and follicular lymphoma (FL), the two most common types of B-cell non-Hodgkin lymphoma, arise in B-cells found in germinal centers of the lymph nodes^1-4^. Mutations in chromatin regulators are frequent in these lymphomas, including recurrent change-of-function missense mutations in the SET domain of *EZH2* at Tyr646, Ala682 or Ala692 (residue numbering per canonical EZH2 isoform Q15910-1, corresponding to Y641/A677 of the frequently referenced Q15910-2 isoform), which are present in approximately 25% of DLBCL and FL cases^5-10^. Covalent EZH2 inhibitors were developed as a precision medicine strategy for lymphomas carrying these mutations, as well as SMARCB1-mutant sarcomas, both of which are dependent on EZH2^11-15^. However, consistent with prior experience using targeted monotherapies, responses to EZH2 inhibitors are typically varied and short-lived, with resistance often emerging within a year^16,17^. While recent work in T-cell lymphoma has identified acquired mutations in *EZH2* that interfere with drug binding, intriguingly, about half of the patients who develop resistance to EZH2 inhibitors have alternative mechanisms of resistance, which remain poorly understood^18^. Resistance to EZH2 inhibitors in SMARCB1-mutant sarcomas has been linked to loss of NSD1, a histone H3K36 methyltransferase^19^, while in B-cell lymphoma, evidence is limited to *in vitro* studies that implicate upregulation of the MAPK pathway in EZH2 inhibitor resistance^20^. An additional challenge is that EZH2 inhibitor therapy has been implicated in secondary malignancies^21,22^. To address these challenges, it will be crucial to determine whether oncogenic *EZH2* mutations create additional exploitable genetic vulnerabilities in EZH2-dependent lymphomas.

EZH2 is the enzymatic component of the Polycomb Repressive Complex 2 (PRC2), which mediates all mono-, di- and tri-methylation of lysine 27 on Histone H3 (H3K27me1/2/3). The core PRC2 complex is composed of EZH2 (or its paralog EZH1), SUZ12 and EED^23-25^. In mammals, PRC2 exists as two distinct subcomplexes, PRC2.1 and PRC2.2^26-29^. PRC2.1 incorporates a single Polycomb-like protein (PHF1, MTF2 or PHF19), together with either EPOP or PALI1/2^30-32^, whereas PRC2.2 incorporates AEBP2 and JARID2^26,33^. Oncogenic missense mutations in *EZH2* in B-cell lymphoma change the substrate preference of the enzyme from H3K27me0/1 to H3K27me2^34^. As a result, the interplay between the wild-type and mutant *EZH2* alleles causes a widespread increase in H3K27me3 levels across the genome, with this repressive mark extending into neighboring promoter and intergenic regions, leading to silencing of entire topologically-associated chromatin domains^35-39^. Notably, this increase in H3K27me3 coincides with a global reduction in H3K27me2, a modification typically enriched at intergenic regions but whose function remains poorly understood^38,40-42^. Although PRC2.2 deposits H3K27me2 during early development and has been linked to transcriptional repression, how H3K27me2 is regulated and the consequences of its dysregulation in EZH2-mutant lymphoma remain unknown^43,44^.

Here, we identify a previously uncharacterized form of PRC2.2, consisting of the long isoform of AEBP2 (AEBP2^L^) but lacking JARID2, which represents a specific dependency in EZH2-mutant B-cell lymphoma. Loss of AEBP2^L^ impairs proliferation specifically in EZH2-mutant B-cell lymphomas, but not in other PRC2-dependent cancers such as malignant rhabdoid tumors. While AEBP2^L^ loss causes global increases in H3K27me3, we demonstrate that the accompanying reduction in H3K27me2, rather than this elevated H3K27me3, drives the proliferation defect. We further establish that the tandem zinc finger domains of AEBP2^L^ are essential for promoting PRC2-mediated H3K27me2 deposition at intergenic regions. Through comparative profiling of cell proliferation rates and H3K27me2 deposition across *MTF2, NSD2* and *AEBP2* knockout lymphoma lines, we define a critical threshold of intergenic H3K27me2 required to sustain cell growth and confer resistance to PRC2 inhibitors. Additionally, we demonstrate that targeting AEBP2 can overcome EZH2 inhibitor resistance in genetically distinct contexts, including (1) resistance driven by the well-established *EZH2-Y111* mutation, and (2) the previously unreported mechanism of resistance by NSD2 loss. Collectively, our findings highlight AEBP2^L^-PRC2.2-dependent maintenance of intergenic H3K27me2 as a new therapeutic vulnerability for these tumors. More broadly, we establish that H2K27me2 functions as a key modulator of sensitivity or resistance to PRC2 inhibitors and that its dysregulation represents a previously underappreciated form of PRC2 dysfunction in cancer.

## RESULTS

### AEBP2 is an EZH2 co-dependency in EZH2-mutant B-cell lymphoma but dispensable for H3K27me3-mediated gene repression

We performed a genome-wide CRISPR-Cas9 screen in the EZH2 Y646F germinal center B-cell lymphoma line WSU-DLCL2 to define essential genes in EZH2-mutant lymphoma (Figure 1A). Consistent with previous work^13,14,45-47^, cells containing single guide RNAs (sgRNAs) targeting core PRC2 components (EZH2, SUZ12 and EED) showed significant dropout in the EZH2-mutant lymphoma cell population over time (Figure 1A). Surprisingly, amongst the sgRNAs targeting PRC2 accessory components, only those targeting *AEBP2* were depleted. To examine whether *AEBP2* dependency is universal in PRC2-dependent cancers, we performed validation assays in SMARCB1-mutant extracranial malignant rhabdoid tumor (eMRT)^15^. To do this, we measured the depletion of GFP-tagged sgRNAs in CRISPR-Cas9 growth-competition assays, targeting core PRC2 components (EED and SUZ12) and PRC2.2 subunits (AEBP2 and JARID2) in EZH2-mutant lymphoma (WSU-DLCL2) and SMARCB1-mutant eMRT (G401) cells (Figure 1B). Strikingly, sgRNAs directed against core PRC2 components caused robust depletion in both cancer types, whereas sgRNAs targeting *AEBP2* selectively depleted in lymphoma cells. Unexpectedly, sgRNAs targeting *JARID2* did not score, suggesting that AEBP2 within PRC2.2, but not JARID2, is required for EZH2-mutant lymphoma cell survival. As an orthogonal validation, we expressed HA-tagged AEBP2^L^, the predominant isoform in these cells, and performed negative selection assays using two independent short hairpin RNAs (shRNAs) targeting the 3’ untranslated region (3’UTR) of endogenous AEBP2, but not the exogenous AEBP2^L^ ORF. HA-AEBP2^L^ expression rescued the proliferation defect in AEBP2-depleted but not EZH2-depleted cells, confirming the specificity of the dependency (Figure S1A-C).

**Figure 1.**
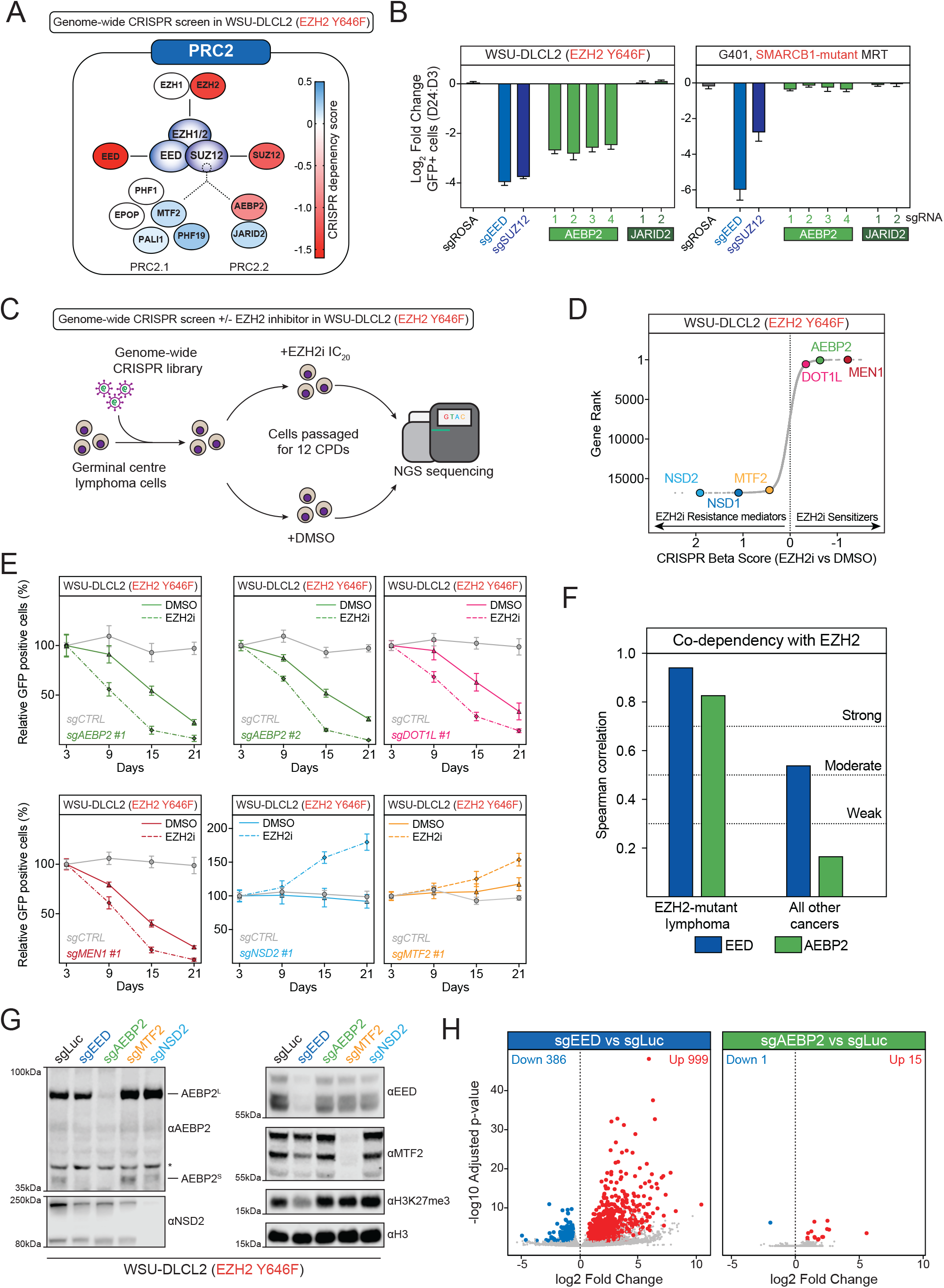
AEBP2 is an EZH2 co-dependency in B-cell lymphoma but dispensable for H3K27me3-mediated gene repression. **A**. Heatmap representation of CRISPR dependency scores for all PRC2 members from genome-wide CRISPR screen in WSU-DLCL2 (EZH2-mutant lymphoma). Screens were performed in 2 biological replicates. **B**. Growth competition assays in Cas9-expressing WSU-DLCL2 (EZH2-mutant lymphoma) and G401 (SMARCB1 mutant malignant rhabdoid tumor) cell lines transduced with sgRNAs targeting the indicated genes (n=3, data represents mean ± SD). *(legend continued on next page)* **C**. Schematic of the genome-wide CRISPR-Cas9 screen performed in germinal center B-cell lymphoma cells. CPD, cell population doublings; EZH2i, EZH2 inhibitor (Tazemetostat). **D**. Genome-wide CRISPR screen data in WSU-DLCL2 (EZH2-mutant lymphoma) cells gene ranked based on sgRNA enrichment in low dose EZH2i treatment (Tazemetostat) relative to DMSO control. Differential Beta-score between EZH2i and DMSO conditions calculated using the MAGeCK algorithm^48^. A negative Beta score indicates sensitizer genes, while a positive Beta score indicates resistor genes. **E**. Growth competition assays in Cas9-expressing WSU-DLCL2 (EZH2-mutant lymphoma) transduced with sgRNAs targeting the indicated genes followed by treatment with low dose EZH2i (Tazemetostat) (n=3, data represents mean ± SD). **F**. Correlation scores for *EZH2* co-dependencies with either *EED* or *AEBP2*, produced using CHRONOS gene-effect scores in the Cancer Dependency Map (https://depmap.org/portal/). Co-dependencies are divided into EZH2-mutant lymphoma (n=6) and all other cancer cell lines (n=995). **G**. Immunoblots of the indicated PRC2 proteins and histone modifications in WSU-DLCL2 (EZH2-mutant lymphoma) cells transduced with the indicated sgRNAs. The * denotes a background band. **H**. Volcano plots of RNA-seq depicting differentially expressed genes in Cas9-expressing WSU-DLCL2 (EZH2-mutant lymphoma) cells transduced with sgRNAs targeting *EED* or *AEBP2*. The number of genes differentially expressed both up (red) and down (blue) are indicated on the plots (n=3 biological replicates; significance cutoff: p < 0.05; minimum 1.5-fold change). **A-H**. Core PRC2 complex is shown in blue, PRC2.1 accessory proteins are shown in orange, and PRC2.2 accessory proteins are shown in green.

Next, to identify genes whose loss sensitizes EZH2-mutant lymphoma cells to the EZH2 inhibitor Tazemetostat, we compared CRISPR Beta scores^48^ between cells treated with DMSO (Control) and those exposed to a low dose (IC_20_) of Tazemetostat (Figure 1C). A negative Beta score indicates a gene whose loss sensitizes cells to EZH2 inhibition, whereas a positive score indicates genes whose loss confers resistance. *AEBP2* ranked 71 out of 16,808 genes, identifying it as a strong sensitizer to EZH2 inhibition, alongside two previously characterized sensitizer genes, *MEN1* (7/16,808)^49^ and *DOT1L* (561/16,808)^50^ (Figure 1D). The only other PRC2 accessory component to score in the screen was the PRC2.1 specific component *MTF2*, which displayed a mild resistance phenotype (16,448/16,808), in contrast to the *AEBP2* phenotype. Additionally, we found that the H3K36me2 methyltransferases NSD1 (16,778/16,808) and NSD2 (16,805/16,808) ranked among the top genes conferring resistance (Figure 1D). Notably, we and others have shown that loss of either *NSD1* or *NSD2* leads to global increases in H3K27me3 levels^19,51-53^. A previous study demonstrated that *NSD1* loss confers resistance to EZH2 inhibitors in MRT by reducing H3K36me2, a histone modification that they reported is required for EZH2 inhibitor-induced gene activation and therapeutic efficacy, thus providing a potential mechanistic basis for the resistance phenotype^19^.

We next validated our CRISPR screen using CRISPR-Cas9 negative selection assays in the presence or absence of Tazemetostat (Figure 1E, S1D). Two independent sgRNAs targeting *AEBP2* sensitized lymphoma cells to Tazemetostat treatment, confirming *AEBP2* as a genetic sensitizer. As expected, *MEN1* and *DOT1L* also produced sensitizing effects (Figure 1E, S1D). In contrast, the loss of *MTF2* and *NSD2* conferred resistance to Tazemetostat, consistent with findings from the genome-wide screen. To evaluate the broader relevance of *AEBP2* as an *EZH2* co-dependency, we analyzed data from the Cancer Dependency Map (https://depmap.org/portal) to compare the strength of *EZH2-AEBP2* co-dependency in EZH2-mutant lymphomas versus other cancer types (Figure 1F). Notably, the co-dependency between *EZH2* and *AEBP2* was comparable to that between *EZH2* and *EED* in EZH2-mutant lymphoma, but much weaker in other cancer types; EED being a core PRC2 component and a proposed therapeutic target in lymphoma^45,47,54^.

To better understand why *AEBP2* is an *EZH2* co-dependency in lymphoma, we first assessed global H3K27me3 levels by immunoblot after CRISPR-mediated depletion of both isoforms of *AEBP2*, using *EED* depletion as a control. As expected, loss of *EED* reduced global H3K27me3 levels, whereas *AEBP2* depletion caused H3K27me3 levels to remain largely unchanged (Figure 1G). Thus, while loss of *EZH2* or *EED* reduces H3K27me3, the fact that *AEBP2* depletion does not indicates that the *AEBP2-EZH2* co-dependency is unlikely to be related to global H3K27me3 levels. We next compared the transcriptional effects of core PRC2 depletion or inhibition with those of *AEBP2* loss, using CRISPR-Cas9, shRNA and pharmacological approaches (Figure 1H, S1E-H). Disruption of core PRC2 components or treatment with Tazemetostat led to widespread transcriptional de-repression, consistent with reduced H3K27me3 levels. In contrast, *AEBP2* depletion caused minimal gene expression changes.

Together, these results show that *AEBP2* is both a genetic dependency and a sensitizer to EZH2 inhibition in EZH2-mutant B-cell lymphoma, yet its loss does not lead to global H3K27me3 reduction or phenocopy activation of genes typically repressed by PRC2.

### *AEBP2* and *NSD2* loss both increase intergenic H3K27me3 while paradoxically conferring opposing sensitivity to EZH2 inhibitors

We next investigated whether the AEBP2 loss could overcome resistance to EZH2 inhibitors. To explore this, we first evaluated loss of AEBP2 in an isogenic EZH2-mutant lymphoma model (Pfeiffer) expressing the EZH2-Y111D/A677G double mutant, which conferred resistance to Tazemetostat (Figure S2A-C). Our results showed that AEBP2 loss, using two independent sgRNAs, effectively blocked proliferation regardless of the presence of the EZH2-Y111D mutation (Figure S2D). We also evaluated NSD2-KO lymphoma cells using cell viability assays, analyzing populations completely lacking the targeted genes (Figure 2A-B). As expected, loss of NSD2 alone conferred resistance to Tazemetostat, whereas AEBP2 loss enhanced sensitivity. Notably, the loss of AEBP2 in NSD2-KO cells could partially overcome Tazemetostat resistance, suggesting that dual targeting of EZH2 and AEBP2 could be an effective strategy to prevent resistance development *in vivo* (Figure 2A).

**Figure 2.**
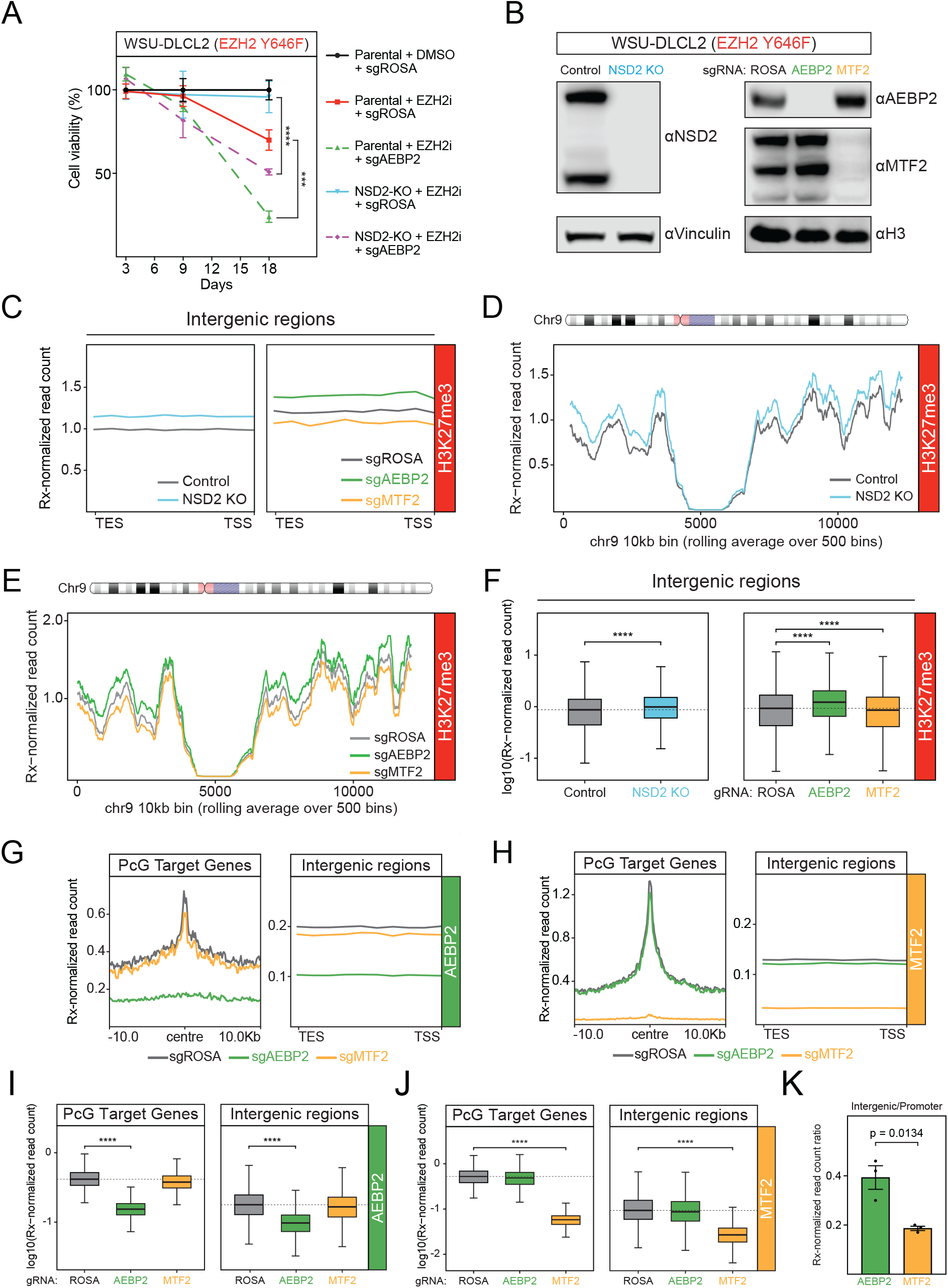
*AEBP2* and *NSD2* loss both increase intergenic H3K27me3 despite conferring opposing sensitivity to EZH2 inhibitors. **A**. Cell viability assay using the CellTiter-Glo® reagent in WSU-DLCL2 lymphoma cells with CRISPR KO of the indicated genes and treated with DMSO or Tazemetostat (n=3, data represents mean ± SD). Significance determined by ordinary one-way ANOVA with Tukey’s multiple comparison test, ***p value ≤ 0.001, ****p value ≤ 0.0001. **B**. Immunoblots of the indicated proteins in WSU-DLCL2 (EZH2-mutant lymphoma) clonal *NSD2* KO and control (sgROSA) isogenic cell lines (left) as well as Cas9-expressing WSU-DLCL2 cells transduced with sgRNAs targeting the indicated PRC2 genes (right). **C**. Average plots showing spike-in normalized H3K27me3 CUT&RUN-Rx enrichment at all intergenic sites in control and *NSD2* KO WSU-DLCL2 (EZH2-mutant lymphoma) cells (left) and WSU-DLCL2 (EZH2-mutant lymphoma) cells transduced with sgRNAs targeting the indicated PRC2 genes (right). TES and TSS indicate Transcription End Site and Transcription Start Site, respectively. *(legend continued on next page)* **D**. Rolling average plots presenting H3K27me3 CUT&RUN-Rx signal across the whole of chromosome 9 in control and *NSD2* KO WSU-DLCL2 (EZH2-mutant lymphoma) cells. Visualized using 10kb genomic windows. **E**. Rolling average plots presenting H3K27me3 CUT&RUN-Rx signal across the whole of chromosome 9 in Cas9-expressing WSU-DLCL2 (EZH2-mutant lymphoma) cells transduced with sgRNAs targeting the indicated PRC2 genes. Visualized using 10kb genomic windows. **F**. Boxplots of H3K27me3 CUT&RUN-Rx signal at intergenic regions in control and *NSD2* KO WSU-DLCL2 (EZH2-mutant lymphoma) cells and in Cas9-expressing WSU-DLCL2 (EZH2-mutant lymphoma) cells transduced with sgRNAs targeting the indicated PRC2 genes. Significance determined using pairwise Wilcoxon rank-sum tests, ****p value ≤ 0.0001. **G**. Average plots showing AEBP2 CUT&RUN-Rx enrichment at genomic windows +/-10kb from PRC2-bound promoters (left) and at all intergenic sites (right) in Cas9-expressing WSU-DLCL2 (EZH2-mutant lymphoma) cells transduced with sgRNAs targeting the indicated PRC2 genes. **H**. Average plots showing MTF2 CUT&RUN-Rx enrichment at genomic windows +/-10kb from PRC2-bound promoters (left) and at all intergenic sites (right) in Cas9-expressing WSU-DLCL2 (EZH2-mutant lymphoma) cells transduced with sgRNAs targeting the indicated PRC2 genes. **I**. Boxplots of AEBP2 CUT&RUN-Rx signal at PRC2-bound promoters (left) and intergenic regions (right) in Cas9-expressing WSU-DLCL2 (EZH2-mutant lymphoma) cells transduced with sgRNAs targeting the indicated PRC2 genes. Significance determined using pairwise Wilcoxon rank-sum tests, ****p value ≤ 0.0001. **J**. Boxplots of MTF2 CUT&RUN-Rx signal at PRC2-bound promoters (left) and intergenic regions (right) in Cas9-expressing WSU-DLCL2 (EZH2-mutant lymphoma) cells transduced with sgRNAs targeting the indicated PRC2 genes. Significance determined using pairwise Wilcoxon rank-sum tests, ****p value ≤ 0.0001. **K**. Ratio of CUT&RUN-Rx signal at intergenic regions compared to PRC2-bound promoter regions for both AEBP2 and MTF2 (n=3, data represents mean ± SEM). Significance determined using unpaired t-test.

Given the starkly opposing consequences of *AEBP2* compared to *NSD2* loss on EZH2 inhibitor response, we hypothesized that the genome-wide profiles of H3K27me3 might differ between the two contexts. Although quantitative western blots showed little changes the global levels of H3K27me3 in both *AEBP2*- and *NSD2*-deficient cells, we reasoned that such bulk measurements might mask qualitative alterations in the distribution of H3K27me3, as we and others first reported for embryonic stem cells (ESCs) lacking accessory proteins^29,55^. We therefore examined the genome-wide distribution of H3K27me3 in EZH2-mutant lymphoma cells with loss of *AEBP2* versus loss of *NSD2*, using *MTF2* loss as a positive control since it is known to reduce H3K27me3 at Polycomb target genes^29,49,55^. To test this, we performed spike-in normalized CUT&RUN (CUT&RUN-Rx) profiling for H3K27me3 and found that loss of *AEBP2* caused a genome-wide increase in H3K27me3, most notably at intergenic regions, as did loss of *NSD2* (Figure 2C-F, S2E-G). In contrast, *MTF2* loss caused a decrease in intergenic H3K27me3 with more pronounced reductions at Polycomb target genes, while paradoxically promoting resistance to EZH2 inhibition. We also compared the effects of shRNA mediated depletion of EZH2 and AEBP2 on H3K27me3 levels and found that, in contrast to loss of AEBP2, EZH2 loss significantly reduces H3K27me3 (Figure S2H-J). Taken together, these findings indicate that changes in H3K27me3 alone do not appear to account for the differential sensitivity of EZH2-mutant B-cell lymphoma cells to loss of *NSD2, MTF2, AEBP2* and *EZH2*. Instead, they suggest AEBP2 has an additional, distinct function beyond modulation of H3K27me3 levels.

To further investigate the role of AEBP2 in EZH2-mutant B-cell lymphoma, we profiled its genome-wide binding using CUT&RUN-Rx, comparing it to MTF2 (Figure 2G-K and S2K-L). We first confirmed the antibody specificity for AEBP2 and MTF2 in the CUT&RUN assays using knockout lymphoma cells (Figure 2B) and found that while both proteins bound Polycomb target gene promoters, they appeared to display distinct preferences (Figure 2G-J). While MTF2 appeared more strongly enriched at promoter-associated PRC2-binding sites, AEBP2, while also binding at Polycomb target promoters, appeared to show relatively greater occupancy at intergenic regions (Figure 2K, S2K-L). This pattern suggested to us that AEBP2 could also function outside of canonical Polycomb target gene promoters, potentially playing a distinct regulatory role at intergenic genomic regions.

### AEBP2 confers its essential function in EZH2-mutant lymphoma in a PRC2 complex lacking JARID2

Although JARID2 is a well-established component of PRC2.2 alongside AEBP2, we were surprised to observe that sgRNAs targeting *JARID2* were not depleted in our genome-wide CRISPR screens of EZH2-mutant lymphoma cells (Figure 1A). Given that only four sgRNAs per gene are included in this genome-wide library, we sought to use our previously reported saturating CRISPR-Cas9 library, tiling 3,599 sgRNAs across all PAM sequences in the exons of genes encoding core and accessory PRC2 proteins, to more rigorously evaluate genetic dependencies (Figure 3A)^56^. As expected, sgRNAs targeting core PRC2 components *EZH2, SUZ12* and *EED* showed strong depletion (Figure 3B). *AEBP2* also scored as a dependency with multiple sgRNAs showing depletion, whereas *JARID2*-targeting sgRNAs did not, definitively ruling out *JARID2* as a dependency. Importantly, we further demonstrated that multiple independent EZH2-mutant B-cell lymphoma cell lines are functionally dependent upon *AEBP2*, but not *JARID2*, using CRISPR-Cas9 negative selection assays (Figure 3C), employing multiple independent sgRNAs, including those specifically targeting the previously described SUZ12-binding region of AEBP2^57^. This provided functional evidence that AEBP2 acts through PRC2, with its role in lymphoma dependent on this interaction.

**Figure 3.**
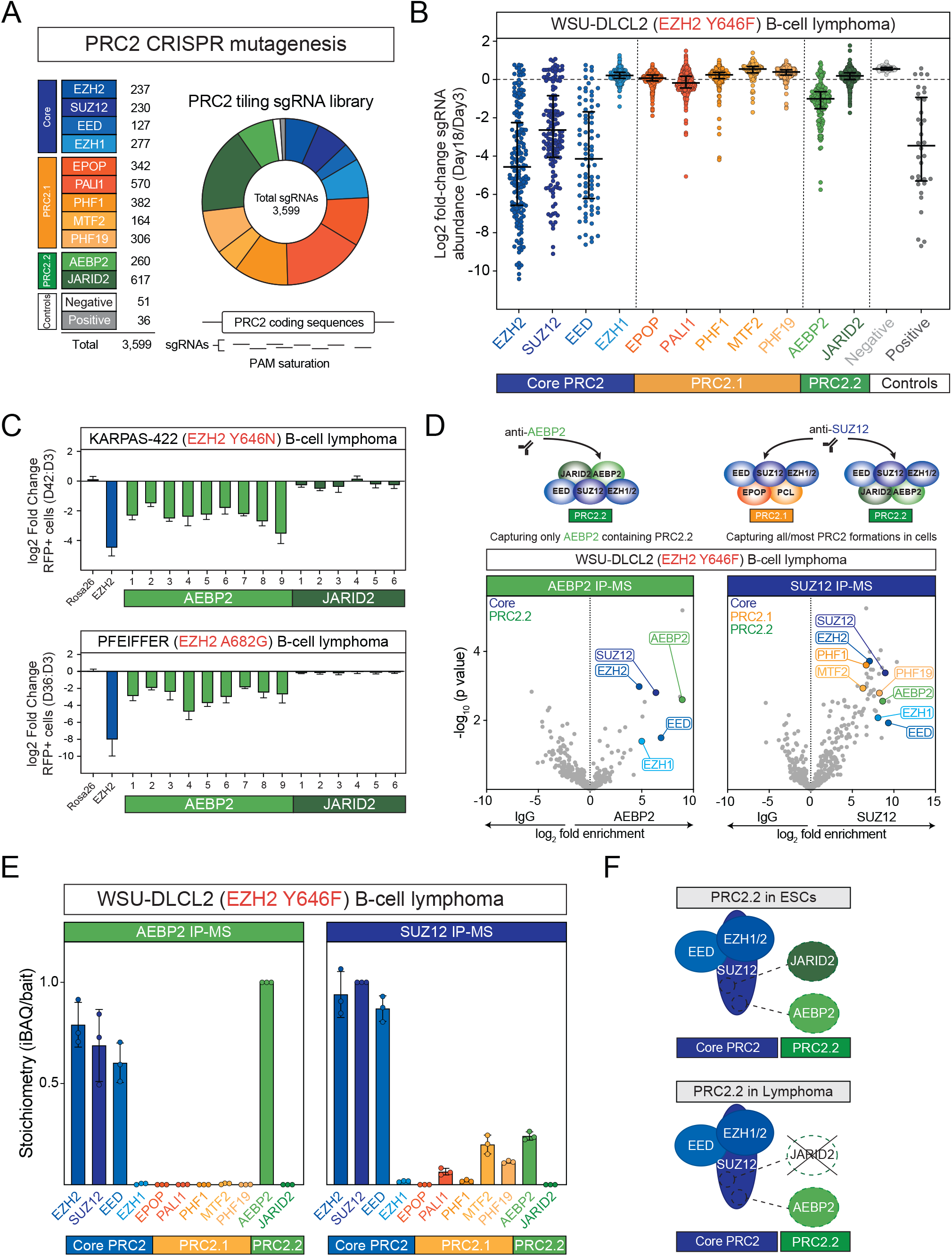
AEBP2 confers its essential function in EZH2-mutant lymphoma in a PRC2 complex lacking JARID2. **A**. Schematic of PRC2-tiling CRISPR library. Indicated are the total number and proportion of sgRNAs targeting each complex member. **B**. PRC2-tiling CRISPR dropout screen in WSU-DLCL2 (EZH2-mutant lymphoma) cells. Each dot represents the Log_2_ Fold Change (mean of n = 2) of individual sgRNAs targeting PRC2 members and controls in pooled CRISPR dropout experiments. The median and interquartile range of all sgRNAs targeting a given gene are indicated. **C**. Growth competition assays in Cas9-expressing Karpas-422 and Pfeiffer (EZH2-mutant lymphoma) cells expressing sgRNAs targeting each of the indicated AEBP2 (left) and JARID2 (right) genes (n = 3, data represent mean ± SD). *(legend continued on next page)* **D**. Schematic of the SUZ12 co-IP-MS strategy to capture all PRC2 complexes, compared with the AEBP2 co-IP-MS strategy to capture only PRC2.2 complexes (top) and volcano plots (bottom) illustrating protein enrichment in endogenous co-immunoprecipitations followed by mass spectrometry of AEBP2 (left) and SUZ12 (right) in WSU-DLCL2 (EZH2-mutant lymphoma) cells (n=3 biological replicates). **E**. Bar plots depicting stoichiometry from the IP-MS of AEBP2 (left) and SUZ12 (right) in WSU-DLCL2 (EZH2-mutant lymphoma) cells (n=3, data represents mean ± SD). **F**. Schematic depictions of core PRC2 complex interacting with PRC2.2 accessory proteins in embryonic stem cells compared to B-cell lymphoma cells. **A-F**. Core PRC2 complex is shown in blue, PRC2.1 accessory proteins are shown in orange, and PRC2.2 accessory proteins are shown in green.

Next, to investigate the composition of PRC2 in EZH2-mutant B-cell lymphoma cells, we performed endogenous co-immunoprecipitation followed by mass spectrometry (IP-MS), using antibodies against AEBP2 or the core PRC2 component SUZ12 (Figure 3D). This enabled us to capture all PRC2 formations as well as those specifically containing AEBP2. As expected, both AEBP2 and SUZ12 co-immunoprecipitated core PRC2 components; however, JARID2 peptides were not enriched in either IP-MS from EZH2-mutant B-cell lymphoma cells (Figure 3D). Quantitative analysis of SUZ12 IP-MS revealed that MTF2 and AEBP2 were the predominant PRC2.1 and PRC2.2 accessory proteins, respectively, in these cells (Figure 3E). Notably, AEBP2 was detected only in complexes with core PRC2, with no evidence of alternative interacting proteins, further supporting the idea that it exerts its functions exclusively through PRC2. The *AEBP2* gene locus encodes two isoforms, AEBP2^L^ (long) and AEBP2^S^ (short), produced via alternative promoters^58-60^. Immunoblotting revealed that AEBP2^L^ is the predominant isoform across a panel of B-cell lymphoma cell lines (Figure S3A). Collectively, these findings reveal a novel PRC2.2 configuration in EZH2-mutant B-cell lymphoma: a complex containing AEBP2^L^ but lacking JARID2, which we term AEBP2^L^-PRC2.2 (Figure 3F), distinct from the canonical JARID2-AEBP2-containing PRC2.2 characterized in mouse embryonic stem cells (ESCs).

### AEBP2^**L**^ **chromatin binding via its zinc fingers is critical for its role in lymphoma**

One possible explanation for the impaired proliferation following *AEBP2* loss could be increased formation of MTF2-PRC2.1 complexes, due to reduced competition for SUZ12 binding^25,33,57^, leading to aberrant accumulation of H3K27me3. If this were the case, only the SUZ12-binding helix (SBH) of AEBP2 would be required for its function, while its zinc finger (ZF) domains would be dispensable. To investigate this, we analyzed sgRNA depletion across *AEBP2, EZH2* (positive control), and *JARID2* (negative control) from our PRC2-tiling CRISPR screen (Figure 3A), mapping sgRNA dropout to specific protein regions and identifying CRISPR-knockout hyper-sensitive (CKHS) sites using the ProTiler^61^ algorithm (Figure 4A). Importantly, for a region to score as a CKHS, the algorithm requires runs of consecutive low-scoring sgRNAs and excludes those with poor-specificity. As expected, the EZH2 SET domain scored as a CKHS region. While two sgRNAs targeting the AEBP2 PRC2-interacting region also showed strong dropout, intriguingly we observed strong depletion in the three ZF domains of AEBP2, indicating their functional importance. In contrast, no hypersensitive regions were detected in JARID2, consistent with its non-essentiality. For comparison, we separately confirmed that *AEBP2, EZH2* and *JARID2* contain missense-intolerant regions in the general population (Figure S4A), and though this does not imply cancer-specific dependency, it indicates that the ZFs of AEBP2 are specifically dependent in EZH2-mutant B-cell lymphoma. We next performed a PRC2-tilling screen in the G401 eMRT cells and, while the EZH2 SET domain scored as expected, the AEBP2 PRC2-interacting region and ZF domains did not (Figure S4B), further indicating that their importance is specific to lymphoma.

**Figure 4.**
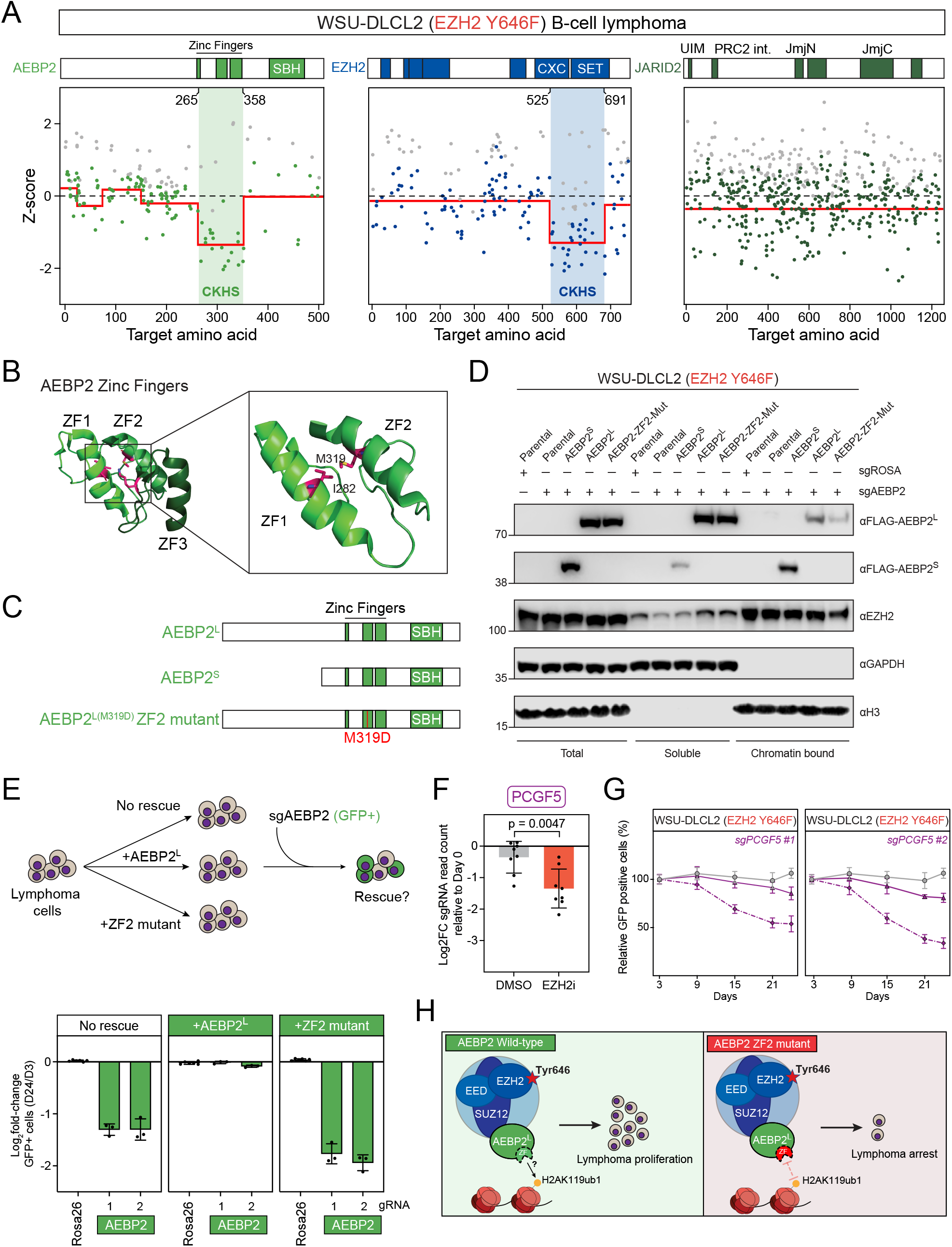
AEBP2 chromatin binding via zinc fingers is critical for its role in lymphoma. **A**. CRISPR tiling screen results of AEBP2, EZH2 and JARID2 in WSU-DLCL2 (EZH2-mutant lymphoma) cells (n = 2 biological replicates). sgRNAs are indicated by dots mapped to the target amino acid. CRISPR-knockout hyper-sensitivity (CKHS) regions are indicated in the shaded regions. The average depletion of sgRNAs targeting a protein segment is indicated by the red line (Z-score). The relevant domains of each protein are indicated. SBH indicates the SUZ12-binding helix domain. **B**. Structures of human AEBP2 zinc finger domains, determined by NMR spectroscopy^62^, with the methionine 319 mutation target indicated. **C**. Diagrammatic representation of the HA/FLAG-tagged AEBP2 long, AEBP2 short, and zinc finger 2 (ZF2) mutant constructs expressed in WSU-DLCL2 (EZH2-mutant lymphoma) cells. SBH indicates the SUZ12-binding helix domain. *(legend continued on next page)* **D**. Immunoblots showing cellular fractionations in WSU-DLCL2 (EZH2-mutant lymphoma) cells expressing the HA/FLAG-tagged AEBP2 long, AEBP2 short, and zinc finger 2 mutant constructs, using the indicated antibodies. Cells expressing sgROSA (control) or sgAEBP2 are indicated. **E**. Schematic representation of AEBP2 rescue experiment (top) and growth competition assays (bottom) in WSU-DLCL2 (EZH2-mutant lymphoma) cells expressing sgRNAs targeting endogenous AEBP2 with or without CRISPR-resistant AEBP2^L^ or AEBP2 ZF2 mutant rescue constructs (n=3, data represent mean ± SD). **F**. Log2 Fold change of individual sgRNAs relative to D0 from either DMSO-treated or Tazemetostat-treated WSU-DLCL2 cells from the genome-wide CRISPR screen (Figure 1C), p value calculated by Mann-Whitney test. **G**. Growth competition assays in Cas9-expressing WSU-DLCL2 (EZH2-mutant lymphoma) transduced with sgRNAs targeting the indicated genes followed by treatment with low dose EZH2i (Tazemetostat) (n=3, data represents mean ± SD). **H**. Model demonstrating the lymphoma cell phenotype in cells expressing wild-type AEBP2^L^ compared to AEBP2 ZF2 mutant. The recurrent Y646X mutation in EZH2-mutant lymphoma cells is indicated with a red star.

To provide higher resolution within the zinc finger CKHS region block of AEBP2, we used a previously validated “smoothen” model, combined with an NMR-determined structure of its ZFs^62^ (Figure 4B, S4C). This approach allowed us to identify methionine 319 within ZF2, the central zinc-finger, as a candidate residue that may be critical for zinc-finger function, allowing us to validate the functional importance of the ZF domains of AEBP2. We hypothesized that substituting methionine 319 with aspartate (M319D), a residue in the ZF2 α-helix that forms to a hydrophobic core with ZF1, would preserve the overall structure of AEBP2 while specifically impairing zinc-finger-mediated chromatin binding, without affecting protein stability or PRC2 association. Ectopic expression of wild-type AEBP2^L^, AEBP2^S^ and the M319D AEBP2^L^ mutant in EZH2-mutant lymphoma cells confirmed stable expression (Figure 4C, S4D), while FLAG co-immunoprecipitations showed that the M319D mutant retained PRC2 binding capability (Figure S4E). However, subcellular fractionation analysis showed a reduced chromatin association for the M319D mutant compared to wild-type AEBP2^L^ (Figure 4D).

We next performed functional rescue assays and found that the M319D mutant was unable to restore proliferation in *AEBP2* knockout lymphoma cells (Figure 4E). Based on the fact that ZF1-2 of AEBP2 have been shown to interact with H2AK119ub1^63^, we hypothesized that the Zinc finger domains of AEBP2 play a role in recognizing intergenic H2AK119ub1, a modification mediated by PCGF3/PCGF5-containing vPRC1 complexes^64,65^. Supporting this hypothesis, we noted that *PCGF5* was one of the most significant sensitizer genes (55/16,808) in our genome-wide CRISPR screen (Figure 4F). To investigate whether the loss of *PCGF5* synergizes with Tazemetostat, we performed CRISPR-Cas9 negative selection assays with and without Tazemetostat treatment (Figure 4G). Two independent sgRNAs targeting *PCGF5* sensitized lymphoma cells to Tazemetostat, confirming it as a genetic sensitizer similar to *AEBP2*.

Taken together, these results demonstrate that the zinc-finger domains of AEBP2 are essential in EZH2-mutant lymphoma, primarily due to their role in mediating chromatin binding. Furthermore, our findings suggest AEBP2-PRC2 functions downstream of PCGF5-vPRC1 as a reader of intergenic H2AK119ub1 (Figure 4H).

### AEBP2^**L**^ **sustains residual H3K27me2 essential for continued proliferation of EZH2-mutant lymphoma**

Next, given that the essential role of AEBP2^L^ in EZH2-mutant lymphoma is independent of its effects on H3K27me3 levels, we hypothesized that its primary function could be to promote PRC2-mediated H3K27me2 deposition at intergenic regions. This idea was supported by our prior work showing that both AEBP2 and H3K27me2 increase at intergenic regions upon *JARID2* loss^29^. Here, in EZH2-mutant B-cell lymphoma cells, which we have shown lack JARID2, AEBP2 appears to be relatively more enriched at intergenic regions than MTF2. Additionally, we previously established that the ZF domains AEBP2^L^, responsible for chromatin-binding, are functionally essential (Figure 4). Since H3K27me3 levels do not correlate with lymphoma proliferation (Figure 2), we reasoned that H3K27me2, the other major PRC2-mediated modification, could be a critical determinant of EZH2 inhibitor sensitivity. Importantly, it has previously been shown that H3K27me2 is already markedly depleted in EZH2-mutant B-cell lymphomas^34,35,42^.

To test this hypothesis, we developed a sensitive ChIP-Rx assay to quantify genomic H3K27me2 in EZH2-mutant lymphomas. We first generated an isogenic lymphoma model by ectopically expressing either wild-type or change-of-function EZH2-Y646F in EZH2-wild-type OCI-LY7 lymphoma cells. Using mouse ESCs for spike-in normalization, this system enabled reliable ChIP-Rx based assessment of the low levels of H3K27me2 abundance and its distribution, which we show is markedly reduced at intergenic regions, in a reciprocal fashion to H3K27me3, in EZH2-mutant lymphoma cells (Figure S5A-C). We next performed both shRNA-mediated depletion and pharmacologic inhibition of EZH2, showing that H3K27me2 levels decrease at intergenic chromatin regions in both cases (Figure S5D–I), supporting the idea that lymphoma cells are sensitive to changes in H3K27me2. Notably, the depletion of AEBP2 led to greater reductions in H3K27me2 when compared with EZH2 depletion (Figure S5D-F). Next, using CUT&RUN-Rx for H3K27me2, we explored the consequences of CRISPR-mediated loss of *AEBP2* or *MTF2*. This demonstrated that loss of *AEBP2* significantly reduced H3K27me2 at intergenic regions, whereas loss of *MTF2* led to increased H3K27me2 at those same regions (Figure 5A-C). Thus, the changes in H3K27me2 levels correspond to the opposite sensitivities of AEBP2- versus MTF2-depleted cells to EZH2 inhibition (Figure 1D-E).

**Figure 5.**
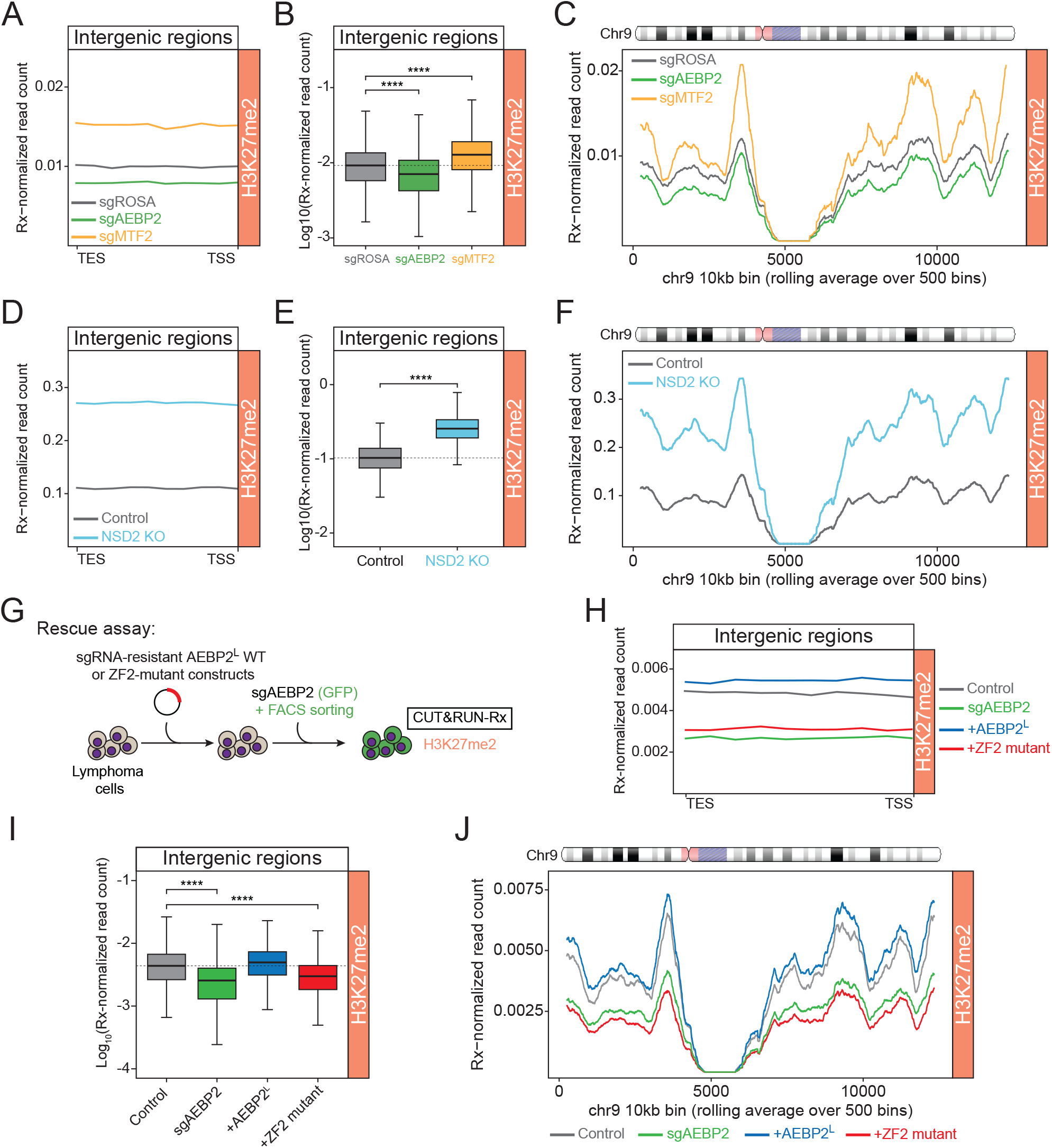
AEBP2 sustains residual H3K27me2 essential for continued proliferation of EZH2-mutant lymphoma. **A**. Average plots showing spike-in normalized H3K27me2 CUT&RUN-Rx enrichment at all intergenic sites in Cas9-expressing WSU-DLCL2 (EZH2-mutant lymphoma) cells transduced with sgRNAs targeting the indicated PRC2 genes. **B**. Boxplots of spike-in normalized H3K27me2 CUT&RUN-Rx signal at all intergenic regions in Cas9-expressing WSU-DLCL2 (EZH2-mutant lymphoma) cells transduced with sgRNAs targeting the indicated PRC2 genes. Significance determined using pairwise Wilcoxon rank-sum tests, ****p value ≤ 0.0001. **C**. Rolling average plots presenting H3K27me2 CUT&RUN-Rx signal across the whole of chromosome 9 in Cas9-expressing WSU-DLCL2 (EZH2-mutant lymphoma) cells transduced with sgRNAs targeting the indicated PRC2 genes. Visualized using 10kb genomic windows. **D**. Average plots showing H3K27me2 CUT&RUN-Rx enrichment at all intergenic sites in control and *NSD2* KO WSU-DLCL2 (EZH2-mutant lymphoma) cells. **E**. Boxplots of CUT&RUN-Rx signal at all intergenic regions in control and *NSD2* KO WSU-DLCL2 (EZH2-mutant lymphoma) cells. Significance determined using pairwise Wilcoxon rank-sum tests, ****p value ≤ 0.0001. **F**. Rolling average plots presenting H3K27me2 CUT&RUN-Rx signal across the whole of chromosome 9 in control and *NSD2* KO WSU-DLCL2 (EZH2-mutant lymphoma) cells. Visualized using 10kb genomic windows. **G**. Schematic of CUT&RUN-Rx experiment in WSU-DLCL2 (EZH2-mutant lymphoma) cells expressing AEBP2 wild-type or mutant constructs and using CRISPR to KO endogenous AEBP2. **H**. Average plots showing H3K27me2 CUT&RUN-Rx enrichment at all intergenic sites in WSU-DLCL2 (EZH2-mutant lymphoma) cells expressing the indicated AEBP2 constructs. *(legend continued on next page)* **I**. Boxplots of H3K27me2 CUT&RUN-Rx signal at all intergenic regions in in WSU-DLCL2 (EZH2-mutant lymphoma) cells expressing the indicated AEBP2 constructs. Significance determined using pairwise Wilcoxon rank-sum tests, ****p value ≤ 0.0001. **J**. Rolling average plots presenting H3K27me2 CUT&RUN-Rx signal across the whole of chromosome 9 in WSU-DLCL2 (EZH2-mutant lymphoma) cells expressing the indicated AEBP2 constructs. Visualized using 10kb genomic windows.

This led us to speculate that H3K27me2 levels could be central to resistance of cells to EZH2 inhibitors. To further investigate this, we profiled *NSD2* knockout cells, which we have shown are more strongly resistant to EZH2 inhibition than *MTF2*-knockout cells (Figure 1). Strikingly, *NSD2* loss led to a more pronounced increase in H3K27me2 at intergenic regions compared to *MTF2* knockout (Figure 5D-F). Notably, the extent of H3K27me2 gain correlated with the degree of resistance to EZH2 inhibitors: *NSD2* knockout conferred strong resistance, while *MTF2* knockout conferred mild resistance, suggesting a dose-dependent relationship between H3K27me2 levels and EZH2 inhibitor resistance.

Finally, we examined whether restoring AEBP2^L^ could rescue H3K27me2 levels in *AEBP2*-deficient cells (Figure 5G). Ectopic expression of AEBP2^L^, but not the chromatin-binding deficient M319D ZF2 mutant, restored H3K27me2 at intergenic regions (Figure 5H-J). This supports a model in which the ZF domains of AEBP2^L^ are required for PRC2 engagement with chromatin at intergenic regions to maintain H3K27me2.

Together, our findings suggest that the characteristically low H3K27me2 levels in EZH2-mutant lymphomas create an Achilles’ heel through a specific dependency on AEBP2-PRC2.2 to maintain this essential modification, with H3K27me2 abundance closely correlating with cellular sensitivity or resistance to EZH2 inhibitors.

## DISCUSSION

EZH2-mutant lymphomas are characterized by globally elevated H3K27me3 and reduced H3K27me2 levels, yet the full functional consequences of this altered PRC2 activity remain unclear. Here, we identify a critical requirement for PRC2-mediated intergenic H3K27me2 in sustaining EZH2-mutant lymphoma proliferation and show that modulation of this mark impacts both sensitivity and resistance to EZH2 inhibitors. We identify a form of PRC2.2 containing AEBP2^L^ but lacking JARID2 that is required for maintaining H3K27me2 at intergenic regions. Our findings uncover a previously unrecognized role for AEBP2-PRC2.2 in mediating H3K27me2 in EZH2-mutant B-cell lymphomas, highlighting it as a potential therapeutic opportunity. More broadly, we establish that H3K27me2 functions as a key modulator of sensitivity or resistance to PRC2 inhibitors, revealing its dysregulation as a previously underappreciated form of PRC2 dysfunction in cancer.

### *AEBP2*^*L*^ joins *MEN1* and *DOT1L* as genes whose loss confer sensitivity to PRC2 inhibitors, albeit via a distinct mechanism

We identify the long isoform of AEBP2 (AEBP2^L^) as a novel EZH2 co-dependency in EZH2-mutant lymphoma, expanding on previous studies that uncovered *MEN1* and *DOT1L* as synthetic genetic vulnerabilities, findings that we also observe in this study^49,50,66^. While inhibition of MEN1 and DOT1L synergize with EZH2 inhibition by enhancing de-repression of PRC2-target genes^49,50,67^, our data suggest that AEBP2 loss functions through a distinct mechanism. Specifically, we show that AEBP2 loss does not reduce H3K27me3 or induce transcriptional activation of PRC2 target genes. Instead, we show that AEBP2^L^ operates within a form of JARID2-free form of PRC2.2 to promote H3K27me2 deposition at intergenic regions - a chromatin modification that is strikingly diminished in EZH2-mutant B-cell lymphomas^38,41,42^.

### AEBP2^L^ is in a JARID2-free form of PRC2.2 that mediates intergenic H3K27me2 deposition

We identify a distinct form of PRC2.2 in EZH2-mutant B-cell lymphoma that contains the long isoform of AEBP2 (AEBP2^L^) but lacks JARID2. Most prior studies of PRC2.2 function have been conducted in mouse ESCs, where both AEBP2 and JARID2 proteins are present as sub-stoichiometric components^29,55,68^. However, JARID2 expression is reduced or lost in some non-embryonic tissue types, including lymphomas^54,69^. Our data thus illuminate a JARID2-independent function of AEBP2–PRC2.2 which is both sufficient and capable of targeting PRC2 to intergenic chromatin to mediate H3K27me2 deposition. While JARID2 recruits PRC2.2 to Polycomb target genes in ESCs, at least in part through the recognition of variant PRC1 (vPRC1) mediated H2AK119ub1^33,70^, AEBP2 can also engage this histone mark via its tandem ZF domains, potentially stabilizing PRC2.2 on chromatin^59,63,71^. However, unlike JARID2 loss, which reduces PRC2 occupancy at target genes, AEBP2 loss does not impair PRC2 binding at Polycomb target genes^33,58^. Instead, we show here that it leads to reductions in intergenic H3K27me2 deposition. Therefore, we propose that AEBP2^L^-PRC2.2 sustains H3K27me2 at intergenic regions through recognition of H2AK119ub1 deposited by variant PRC1 complexes containing PCGF3, or PCGF5^64,71^. Supporting this, we demonstrated in ESCs that AEBP2 is relatively more bound at intergenic chromatin compared to MTF2 and JARID2, and that JARID2 loss results in increased AEBP2-PRC2.2 binding and enhanced H3K27me2 at these regions^29^. Furthermore, we propose that AEBP2^L^-PRC2.2 could act downstream of PCGF5-vPRC1, which functions to mediate intergenic H2AK119ub1^64,65^. Supporting this, we observed PCGF5 in the top 60 sensitizer genes in our genome-wide CRISPR screen and validated that its loss sensitizes lymphoma cells to Tazemetostat treatment. These findings highlight a potential strategy for overcoming resistance to EZH2 inhibitors by disrupting the PCGF5-vPRC1-AEBP2-PRC2.2 axis.

### The potential function of H3K27me2 in B-cell lymphoma

To our knowledge, this is the first study to link a cancer specific genetic vulnerability to aberrantly low H3K27me2 levels and to propose that this may be exploited in cancer therapeutics. We reveal that intergenic H3K27me2 is essential in DLBCL, and that reductions below a critical level impair cellular proliferation, despite no detectable changes in gene expression. In non-cancer cells, H3K27me2 is the most abundant PRC2-mediated histone post translational modification and is mostly located at intergenic chromatin, but its function remains unclear^44,72^. H3K27me2 deposition has been reported to accumulate during early development and *Jarid2* KO leads to increases in H3K27ac at enhancers, coupled with gene activation^43^. However, we did not observe gene activation following the targeted loss of PRC2.2 in EZH2-mutant B-cell lymphoma, via knockout of AEBP2. Given the lack of gene expression changes in *AEBP2* KO, we speculate that H3K27me2 may alternatively function to maintain 3D chromatin structure, genome stability, or impact DNA replication rate or centromere function. Alternatively, it may facilitate recruitment of other chromatin regulators, such as the EHMT1/EHMT2 complex or the H3K27me2-reader NuRD complex complex^73-75^.

To date, the impact of reduced H3K27me2 and its relationship to H3K27me3 has not been examined in EZH2-mutant B-cell lymphomas. While EZH2 inhibitors target both H3K27 methylation states, their putative mechanism of efficacy in B-cell lymphoma is thought to be related to their reduction of H3K27me3 specifically^14,76^. A previously underappreciated aspect of their efficacy may be their further reduction of H3K27me2, which is already diminished due to the *EZH2* mutation^13,14^. Supporting this, we show that *AEBP2* KO synergizes with EZH2 inhibition likely resulting in additional reductions in H3K27me2. We have shown in ESCs that H3K27me2 and H3K27ac are deposited in a reciprocal fashion, with increasing H3K27ac levels also affecting chromatin compaction^77^. Therefore, it is possible that reductions in H3K27me2, due to *AEBP2* loss, may affect chromatin compaction.

### Altered H3K27me2 in cancers as an underappreciated modulator of resistance or sensitivity to EZH2 inhibitors

In contrast to *AEBP2, MEN1* and *DOT1L, NSD1* is a gene whose loss confers resistance to EZH2 inhibitors^19^. Here, we identified *NSD1* and *NSD2* as genes whose loss confer resistance to EZH2 inhibitors in EZH2-mutant B-cell lymphoma, an effect we establish is due to increased intergenic H3K27me2, thus abrogating the efficacy of EZH2 inhibitors. Intriguingly, characterizing H3K27me2 levels in SWI/SNF-mutant cancers with and without NSD1 and with and without EZH2 inhibitors could provide a possible mechanistic link between these two divergent cancer types. We demonstrate that AEBP2^L^-PRC2.2 sustains minimal essential intergenic H3K27me2 in a cancer context where H3K27me2 is markedly scarce, demonstrating also that MTF2-PRC2.1 lacks this function.

Although systematic evaluations of genome-wide H3K27me2 levels in cancers are lacking, other cancers may possess low H3K27me2 and potentially be vulnerable to AEBP2^L^-PRC2.2 targeting. It is known that the NSD family of enzymes inhibit the deposition of H3K27 methylations by PRC2^53,78^, and so cancers with NSD1/2/3 gain-of-function would potentially harbor low H3K27me2 levels, and could therefore be sensitive to loss of AEBP2^79,80^. In contrast, given the antagonistic role between PRC2 and NSD proteins, cancers with loss-of-function *NSD1/2* mutations or containing the H3K36M oncohistone, would be predicted to have reduced H3K36me2 and consequent elevated H3K27me2 levels, thereby rendering them less sensitive to PRC2 inhibitors^53,81-83^. Therefore, we suggest assessing H3K27me2 levels across a broader range of cancers to assess its relevance as a predictive marker of vulnerability to AEBP2 loss and EZH2 inhibition. More broadly, we propose grouping cancers by dysregulated histone modification states rather than specific mutations, to expand the therapeutic scope of targeted chromatin-based drugs.

### AEBP2^L^ as a potential therapeutic target

The biology of AEBP2^L^-PRC2-mediated H3K27me2 offers an interesting potential alternative to targeting the enzymatic activity of EZH2, which faces the challenge of acquired resistance and risks of secondary malignancies^11,18,21^. Given that loss-of-function mutations in AEBP2 are rarely observed in cancers, whereas inactivating mutations/loss of core PRC2 components are well-described^84-86^, targeting the biology of AEBP2^L^ may be an attractive alternative strategy to EZH2 inhibition in EZH2-mutant B-cell lymphoma^21,86-89^. Although AEBP2^L^ lacks a tractable enzymatic domain, targeting its ZF domains^90^ or its interaction with SUZ12 may be feasible. Indeed if AEBP2-binding ligands could be developed it may be targeted using PROTAC-mediated degradation, an emerging therapeutic modality in several cancer types^91-93^. Further work to elucidate the potential upstream function of PCGF3/5-vPRC1-mediated intergenic deposition of H2AK119ub1 could open new avenues for targeting the biology of AEBP2^L^ in lymphoma.

## EXPERIMENTAL MODEL AND SUBJECT DETAILS

### Cell Culture and Lentiviral Production

All cell lines were grown in a humidified incubator at 37°C, 5% CO_2_. Human lymphoma cell lines HT, OCI-LY18, Pfeiffer, WSU-DLCL2, were cultured in suspension in Roswell Park Memorial Institute medium - RPMI 1640 GlutaMAX (Gibco) supplemented with 10% heat-inactivated fetal bovine serum (FBS) (Gibco) and 100U/ml penicillin/streptomycin (Gibco). KARPAS-422 was cultured as above with 20% heat-inactivated FBS. Human lymphoma cell lines OCI-LY1 and OCI-LY7 were cultured in Isocove Modified Dulbecco Media - IMDM (Gibco) supplemented with 20% heat-inactivated FBS and 100U/ml penicillin/streptomycin. Cells were passaged by splitting 8:1 – 12:1 every 2-3 days. HEK293T and G401 (human malignant rhabdoid tumor) cell lines were cultured in Dulbecco’s Modified Eagle Medium - DMEM high glucose (Sigma) supplemented with 10% FBS and 100U/ml pencillin/streptomycin. These cells were grown as monolayers and passaged every 2-3 days using 0.25% Trypsin-EDTA (Gibco). Mouse embryonic stem cells (mESCs) were maintained on 0.1% gelatin-coated culture dishes and passaged every 2 days using 0.25% Trypsin-EDTA. mESCs were cultured under 2i/LIF conditions in Glasgow Minimum Essential Medium (GMEM) (Sigma) supplemented with 20% heat-inactivated FBS, 100u/ml penicillin/streptomycin, 50μM β-mercaptoethanol (Sigma), 1:100 GlutaMAX (Gibco), 1:100 non-essential amino acids (Gibco), 1mM sodium pyruvate (Gibco), 1:500 homemade Leukemia Inhibitory Factor (LIF), 3μM GSK inhibitor CHIRON9902 (Millipore) and 1μM MEK inhibitor PD0325901 (Millipore).

Lentiviral particles were produced using a second-generation lentiviral system in HEK293T cells. HEK293T cells were transfected with a lentiviral expression vector (sgRNA/shRNA/cDNA) or a pooled sgRNA library together with a viral envelope (VSVG) and packaging vector (PAX8) using polyethylenimine (PEI) (Polysciences) following a standard transfection protocol Viral supernatant was harvested 72hr post-transfection, sterile filtered through a 0.45μm polyethersulfone (PES) filter and stored at -80°C.

### Immunoblotting

Cell pellets of human lymphoma cells were lysed in either high salt buffer (50mM Tris-HCl pH 7.2, 1mM EDTA pH 7.4, 300mM NaCl) or RIPA buffer (25 mM Tris-HCl. pH 7.6, 150mM NaCl, 1% NP-40, 1% Sodium Deoxycholate, 0.1% SDS) supplemented with protease inhibitors (aprotinin 2μg/ml, leupeptin 1μg/ml, PMSF 1mM). Protein lysates were sonicated using a Sonifier SFX150 (Branson) or Bioruptor Pico (Diagenode), followed by rotation for 20 minute rotation at 4°C and clarification by centrifugation at 14000rpm for 20 minutes. Protein quantification was performed using the Bradford assay. Denatured protein lysates were separated using 4-12% gradient Bolt Bis-Tris Plus (Invitrogen) gel or 4-12% gradient NuPAGE Bis-Tris (Invitrogen) gel and transferred to nitrocellulose (0.2μm, Amersham) or PVDF (0.45μm, Immobilon-FL) membranes respectively. Membranes were probed with primary and secondary antibodies followed by imaging using the Odyssey Fc Imager (LI-COR).

### Cellular Fractionation

Cellular fractionation of B-cell lymphoma cells was performed as described previously^29^. Resulting lysates were then immunoblotted as described above.

### Endogenous co-immunoprecipitation

Human lymphoma cells were harvested by centrifugation, washed twice with ice cold PBS, and resuspended in Buffer A (25mM HEPES pH 7.6, 5mM MgCl2, 25mM KCl, 0.05mM EDTA, 10% (v/v) glycerol, 0.1% NP-40, 1mM DTT, combined with protease inhibitors) and incubated with rotation at 4°C for 10 minutes. Cell nuclei were pelleted by centrifugation at 500rcf at 4°C for 10 minutes. Nuclei were lysed in Buffer C (10mM HEPES pH 7.6, 3mM MgCl2, 100mM KCl, 0.5mM EDTA, 10% (v/v) glycerol, 1mM DTT, combined with protease inhibitors). Then, 300mM (NH_4_)_2_SO_4_ dissolved in Buffer C was added to the samples which were rotated at 4°C for 10 minutes. Samples were ultra-centrifuged at 350,000rcf for 15 minutes at 4°C using an SW 55 Ti swinging-bucket rotor in a Beckman Coulter Optima L-100XP. The supernatant was collected and nuclear extracts were precipitated by adding 300mg (NH_4_)_2_SO_4_ for each 1ml of sample. The samples were incubated on ice for 20 minutes before repeating ultracentrifugation using the same parameters as above. Nuclear pellets were resuspended in IP buffer (300mM NaCl, 50mM Tris-HCl pH 7.5, 1mM EDTA, 1% (v/v) Triton-X100, 1mM DTT, combined with protease inhibitors) and protein quantification and sample normalization was carried out using the Bradford assay. For each co-IP, 1μl Benzonase Nuclease (Sigma-Aldrich) and 1μg of the respective antibody was added to 1mg of nuclear protein. Samples were rotated at 4°C overnight. The next day washed Protein G Dynabeads (Invitrogen) were added to each sample followed by rotation at 4°C for 2 hours. Beads were then washed four times with IP buffer followed by two cold PBS washes. Beads were stored at -80°C until further processing.

### Mass spectrometry

Beads from the co-immunoprecipitation were washed with phosphate buffered saline and 8M urea was added to a volume of 200μl. Following this, 5mM DTT was added to the beads and incubated at 60°C for 30 minutes. Then, 9mM iodoacetamide solution was added to the beads and incubated in the dark at room temperature for 30 minutes. 2.2mM CaCl_2_ was added to the beads followed by addition of 12.8mM ammonium bicarbonate pH 8. Following this, 1μg of sequencing grade trypsin (Sigma-Aldrich) was added to each reaction and incubated at 37°C overnight on a thermomixer at 350rpm. The next morning, 1% trifluoroacetic acid was added to stop the reaction. The tryptic digests were then desalted using ZipTips (Millipore) according to the manufacturer’s protocol. Peptides (500-1000ng) were loaded onto Evotips as per manufacturer’s instructions (Evosep). Briefly, Evotips were activated by soaking them in isopropanol, primed with 20 µL buffer B (ACN, 0.1% FA) by centrifugation for 1 min at 700 g. Tips were soaked in isopropanol and equilibrated with 20 µL buffer A (MS grade water, 0.1% FA) by centrifugation. Another 20 µL buffer A was loaded onto the tips and the samples were added on top of that. Tips were centrifuged and washed with 20 µl buffer A followed by overlaying the C18 material in the tips with 100 µL buffer A and a short 20 s spin.

The samples were analyzed by the Mass Spectrometry Resource (MSR) in University College Dublin on a Bruker TimsTOF Pro mass spectrometer connected to a Evosep One chromatography system. Peptides were separated on an 8 cm analytical C18 column (Evosep, 3 µm beads, 100 µm ID) using the pre-set 30 samples per day gradient on the Evosep one. The Bruker TimsTOF Pro mass spectrometer was operated in positive ion polarity with TIMS (Trapped Ion Mobility Spectrometry) and PASEF (Parallel Accumulation Serial Fragmentation) modes enabled. The accumulation and ramp times for the TIMS were both set to 100 ms., with an ion mobility (1/k0) range from 0.62 to 1.46 Vs/cm. Spectra were recorded in the mass range from 100 to 1,700 m/z. The precursor (MS) Intensity Threshold was set to 2,500 and the precursor Target Intensity set to 20,000. Each PASEF cycle consisted of one MS ramp for precursor detection followed by 10 PASEF MS/MS ramps, with a total cycle time of 1.16 s.

### Design and synthesis of human PRC2 sgRNA library

The PRC2 complex member gene tiling CRISPR library was designed and synthesized as described previously^56^.

### PRC2-tiling library CRISPR-Cas9 screen

The PRC2 sgRNA lentivirus library pool was generated using an ipUSEPR plasmid - a vector expressing an sgRNA together with a puromycin-resistant gene (Puro) and a RFP fluorescent protein tag. Human lymphoma cell line WSU-DLCL2 was transduced with this lentiviral library at 1000x coverage (1000 cells per sgRNA) and a multiplicity of infection of 10%. For this library of ∼4000 sgRNAs, to ensure coverage of 1000 cells containing each sgRNA, with the MOI of 10%, 40 million cells were infected with the PRC2 sgRNA library and puromycin added at 24 hours for library selection. Cell pellets were harvested at Day 3 post-transduction (early time-point) and Day 18 post-transduction (late timepoint) with two biological replicates per timepoint. Genomic DNA was extracted using the HotSHOT protocol. Cell pellets were lysed in 250μl HotA buffer (0.25mM EDTA, 25mM NaOH: final pH=12) and boiled at 95°C for 30 minutes. DNA was then fragmented by sonication for 15 seconds. 500μl of HotB buffer (40nM Tris-HCl pH 6.8) was added, after which samples were gently vortexed and centrifuged. The purified DNA was then resuspended in DNase/RNase free sigma H_2_O. The purified DNA was then amplified by PCR. Following amplification, PCR products were pooled, run on a 2% agarose gel with excision and purification of the 150bp product band which was subsequently processed for sequencing.

### Genome-wide CRISPR-Cas9 screen

Human lymphoma cell lines (WSU-DLCL2 and HT) with stable Cas9 expression were transduced with lentivirus containing the Brunello genome-wide library of sgRNAs with a coverage of 500x (500 cells per sgRNA) and a low multiplicity of infection (MOI) of 0.15 as previously described^100,115^. During transduction, media was supplemented with 10μg/ml polybrene to improve transduction efficiency. Approximately 24 hours after transduction, cells were selected in 2μg/ml puromycin for 48 hours. At this timepoint (T0), for each cell line, two aliquots of cells were collected to take forward for serial passaging. To one aliquot, the EZH2 inhibitor tazemetostat was added at a concentration of ∼IC20 dose (25nM) and to the other aliquot 0.01% (v/v) DMSO was added. At T0 an aliquot of cells was harvested for genomic DNA extraction. Cells were counted and passaged forward maintaining a 500x coverage of each sgRNA every 3 days. Cells were cultured in tazemetostat or DMSO for 24 days in total, at which point (T24) cells in both conditions were harvested and genomic DNA extracted with the NucleoSpin Blood L (Macherey-Nagel) kit following the manufacturer’s instructions. PCR inhibitors were removed with the OneStep PCR Inhibitor Removal (Zymo Research) kit prior to PCR amplification following the manufacturer’s instructions. The barcoded sgRNA sequences were PCR amplified using the following recipe and PCR steps: DreamTaq Hot Start Green (50μl), P5 index primer 100μM (0.5μl), P7 index primer 100μM (0.5μl) at 95°C for 60 seconds, 28 cycles of 95°C for 30s, 53°C for 30s, 72°C for 30s; then 72°C for 10min. After PCR amplification, DNA fragments were purified using the GeneJET PCR Purification Kit (Thermo Scientific) as per the manufacturer’s instructions. PCR products were then separated by a 2% agarose gel electrophoresis, gel purified using the QIAquick Gel Extraction Kit (Qiagen) and isopropanol precipitated. Size distributions of the PCR products were determined using the 4200 TapeStation System (Agilent). Pooled libraries were diluted and processed for 75-bp single-end sequencing using an Illumina NextSeq 500.

### Negative selection assays

sgRNAs targeting regions of proteins of interest highlighted by the two CRISPR screens were cloned into plasmids containing either a GFP (LRG2.1) or RFP (ipUSEPR) reporter. Lentivirus was produced for these sgRNAs and transduced in Cas9-expressing human cell lines at a target MOI of 0.3. Lentiviral media was then removed, fresh media applied and the cells were cultured for a period of 21-45 days. The cells were split every 3 days at an appropriate ratio with GFP abundance quantified every 3 days using the Guava easyCyte HT system 12 and analyzed using FlowJo software. For shRNA negative selection assays, shRNAs targeting genes of interest were cloned into the GFP expressing plasmid SGEP and human lymphoma cells lines were transduced with lentivirus followed by puromycin selection. GFP positive cells were monitored as per the sgRNA-based negative selection assays above. Sequences of these sgRNAs and shRNAs can be found in Table S1.

### Generation of AEBP2 wild-type and mutant constructs

The ectopic expression of the wild-type and mutant hAEBP2 constructs was performed using in-house lentiviral expression vectors derived from PLENTI EF1A FLAG/HA^31^. Human AEBP2-L (NM_001114176.1) ORF was available in the gateway cloning-compatible entry vector PCR8 (pCR8/GW/TOPO cloning – Invitrogen). Zinc-finger mutant hAEBP2 ORFs were generated by Q5 Site-Directed Mutagenesis (NEB) as per the manufacturer’s protocol and sequences were confirmed using Sanger sequencing. Expression clones were then generated using Gateway LR Clonase II (ThermoFisher).

### shRNA generation for targeted RNA interference

98-mer DNA oligonucleotides were designed using the Broad Institute RNAi Consortium GPP Portal and based on previously published shRNA oligonucleotides^102^. The oligonucleotide was diluted in DNAse, RNAse-free H_2_0 to 0.05ng/μl. As per miRE cloning, overhangs with restriction sites for Xho(5’) and EcoRI(3’) were added by PCR to the oligonucleotide as per Table 2.3 and run on a thermocycler.

### CUT&RUN-Rx

For each CUT&RUN-Rx, human lymphoma cells were combined with mESC (1:10 cell ratio), which served as a 10% exogenous genome “spike-in” for normalization during downstream bioinformatic analysis. Cells were fixed with 0.1% formaldehyde for 1 minute, then quenched with 125mM glycine for 5 minutes and washed with PBS at room temperature. Nuclei were extracted by incubating samples in ice cold nuclear extraction buffer (20mM HEPES pH 7.5, 10mM KCl, 0.1% Triton X-100, 20% Glycerol, 1x protease inhibitor cocktail and 0.5mM spermidine) for 10 minutes. Extracted nuclei were centrifuged at 600rcf and 4°C and resuspended in cold nuclear extraction buffer (100μl per sample). The CUT&RUN was performed as per Epicypher CUT&RUN protocol v1.5.1. CUT&RUN DNA was purified using the Monarch DNA Clean-up kit (New England Biolabs) using a 5:1 binding buffer: sample ratio and following the manufacturer’s instructions.

### CUT&RUN-Rx library preparation

The standard protocol of the NEBNext® Ultra™ II DNA Library prep kit (New England Biolabs) was used for library preparation of the CUT&RUN-Rx samples. Of note, PCR enrichment of adaptor-ligated DNA utilized a total of 14 cycles of denaturation at 98°C for 15 seconds and annealing/short extension at 60°C for 10 seconds. The purified PCR product was quantified using a Qubit fluorometer and fragment size distribution was analyzed on an Agilent Tapestation 4200 using D1000 ScreenTape assay (Agilent). Pooled libraries were diluted and processed for 150-bp paired-end sequencing externally using Novogene.

### Quantitative Chromatin Immunoprecipitation (ChIP-Rx)

Human lymphoma cells were counted and fixed with 1% formaldehyde for 10 minutes, quenched with 125mM glycine for 5 minutes and washed twice with room temperature PBS. Crosslinked cells were lysed in 6ml SDS lysis buffer (10mM NaCl, 50mM Tris pH 8.1, 5mM EDTA pH 8.0, 0.5% SDS, combined with protease inhibitors). Chromatin was pelleted by centrifugation at 1200rpm for 6 minutes at room temperature and resuspended in 1ml of ChIP buffer (2:1 dilution of SDS lysis buffer:Triton dilution buffer [100mM Tris pH 8.6, 100mM NaCl, 5mM EDTA pH8.0, 0.002% NaN_3_, 5% Triton X-100, combined with protease inhibitors]). Chromatin was sonicated to fragments of 200-800bp (Branson Sonifier SFX150 - 1 second on, 4 seconds off at 50% amplitude for total sonication time of 4 minutes). Sonication efficiency was confirmed by agarose gel electrophoresis. Chromatin was quantified using a Qubit dsDNA High Sensitivity Assay Kit (ThermoFisher Q32854) and E14 wild-type mouse embryonic stem cell spike-in chromatin was added to well-sonicated human lymphoma cell line chromatin at a ratio of 1:9, as previously described^116^, prior to overnight incubation with antibody whilst rotating at 4°C. Following overnight incubation, 40μl Protein G Dynabeads (Invitrogen) were added to each sample and incubated for 2 hours at 4°C. Beads were washed three times in mixed micelle wash buffer (150mM NaCl, 20mM Tris pH 8.1, 5mM EDTA pH 8.0, 5.2% sucrose, 1% Triton X-100, 0.2% SDS), twice with Buffer 500 (0.1% sodium deoxycholate, 1mM EDTA pH 8.0, 50mM HEPES pH 7.5, 1% Triton X-100), twice with LiCl detergent wash buffer (0.5% sodium deoxycholate, 1mM EDTA pH 8.0, 250mM LiCl, 0.5% NP-40, 10mM Tris pH 8.0) and once with TE (10mM Tris pH 8.0, 1mM EDTA pH 8.0). Samples were eluted from magnetic beads with elution buffer (0.1M NaHCO_3_, 1% SDS) while shaking for 1 hour at 65°C. The next day, samples were treated with RNaseA (ThermoFisher) at 37°C for 1 hour and Proteinase K (Sigma) at 55°C for 2 hours prior to column-based DNA fragment purification (Qiagen or ThermoFisher). ChIP enrichment was established by qPCR using the SYBR Green I detection system on an Applied Biosystems Quant Studio 3 platform.

### ChIP-Rx library preparation

Purified ChIP DNA was quantified using the Qubit dsDNA High Sensitivity Assay Kit (ThermoFisher). ChIP-Rx sequencing libraries were prepared using the NEBNext Ultra II DNA Library Kit for Illumina (NEB E7645) and NEBNext Multiplex Oligos for Illumina as per the manufacturer’s protocol. Input DNA was standardized per ChIP antibody (range 1-50ng). Following adaptor ligation, DNA was PCR amplified for 4-8 cycles depending on input DNA amount. PCR-amplified DNA was purified using AMPure XP beads (Beckman Coulter A63881) at room temperature. DNA library quality was assessed with a High Sensitivity D1000 Screen Tape (Agilent). The libraries were then used for cluster generation and sequencing using an Illumina NextSeq500, with single end 75bp read length.

### RNA-seq

Total RNA was isolated from WSU-DLCL2 (EZH2-mutant lymphoma) cells 7 days after CRISPR-Cas9 sgRNA-mediated knock-out of *Luciferase* (control), *EED*, and *AEBP2* using the RNeasy Mini Kit (Qiagen) according to the manufacturer’s instructions. RNA-seq libraries were prepared by Novogene as follows. mRNA was purified from total RNA using poly-T oligo-attached magnetic beads. Following fragmentation, first strand of cDNA was synthesized using random hexamers, followed by second strand synthesis using dTTP. cDNA was subject to end repair, A-tailing, adapter ligation before being size selected and amplified for final library generation. Library was assessed by Qubit, Bioanalyser and q-rtPCR for quality control and quantification. Libraries were sequenced paired end to an approximate read depth of 55 million paired end reads on Illumina NovaSeqX plus platform.

### QuantSeq 3’ mRNA sequencing for RNA quantification

RNA was purified from 5 million human lymphoma cells per condition and per replicate as detailed in section 2.3.1 using the RNeasy kit (Qiagen). cDNA was generated from this RNA for quantification by sequencing using the QuantSeq 3’ mRNA-Seq Library Prep FWD Kit for Illumina (Lexogen) using 500ng total RNA as input. The optimal number of PCR cycles for endpoint PCR was determined by qPCR assay using the PCR Add-on Kit for Illumina (Lexogen Cat. No. 020). Individual i7 indexing primers were used for PCR amplificaton. The quality of DNA libraries was examined on a High Sensitivity D1000 Screen Tape (Agilent). The libraries were subsequently used for cluster generation and sequenced using an Illumina NextSeq500, with single end 75bp read length.

### Analysis

#### CRISPR screen data analysis

The PRC2-tiling screen was analyzed as described previously^56^. The MAGeCK algorithm was used to calculate CRISPR beta scores for the genome-wide CRISPR screen using the mle module, comparing sgRNA abundance of gDNA from the end of the screen in DMSO and Tazemetostat conditions, with the original plasmid pool^48^. Common essential genes were defined using the Achilles 19Q3 list of genes and were removed before generating plots using Prism 10 for MacOS (GraphPad).

#### Mass spectrometry data analysis

Bruker mass spectrometric data from the Tims-Tof was processed using the MaxQuant (version 2.0.3.0) incorporating the Andromeda search engine. To identify peptides and proteins, MS/MS spectra were matched against a *homo sapiens* uniprot database containing 82,427 entries (release 2023_03). All searches were performed using the default setting of MaxQuant, with trypsin as the specified enzyme allowing two missed cleavages and a false discovery rate of 1% on the peptide and protein level. The database searches were performed with carbidomethyl as a fixed modification and acetylation (protein N terminus) and oxidation (M) as variable modifications. For the generation of label free quantitative (LFQ) ion intensities for protein profiles, signals of corresponding peptides in different nano-HPLC MS/MS runs were matched by MaxQuant in a maximum time window of 1 min. The ‘proteinGroups.txt’ output file was filtered for contaminants and reverse hits using Perseus version

2.0.10.0. Volcano plots were generated using Perseus version 2.0.10.0. For stoichiometric analysis, iBAQ values for IgG control were first subtracted from each experimental replicate. The subsequent iBAQ value of each identified protein was normalised to the respective bait protein in each IP. The stoichiometry values were averaged across replicates, reported as the mean. Plots were formatted using GraphPad PRISM version 10.5.0.

#### CUT&RUN-Rx data analysis

Firstly, adapters from paired-end sequencing reads were trimmed using fastp^107^. In order to perform alignments with exogenous spike-in mouse reference genome, a human-mouse hybrid genome was created by appending the mouse mm10 genome build to the hg38 human genome build prior to indexing. Reads were aligned to the hybrid genome using Bowtie2^103^ in the -very sensitive mode. SAMtools^105^ was used to keep reads with a mapping quality >2, separate the mouse and human reads, remove duplicate reads, and convert SAM to BAM files. The normalization factor generated by the spike-in for each antibody was calculated using the formula for normalized reference-adjusted reads per million (RRPM) (1 per million spike-in reads). The bamCoverage module from the deepTools2^108^ suite was used to generate RRPM normalized bigwig files for genome browser visualization. Average plots were generated using computeMatrix from the deepTools2 suite, while boxplots were generated using ggplot2 in R, after producing matrices using the deepTools2 suite multiBigwigSummary. Regions that overlapped with a custom hg38 blacklist were excluded from analyses.

#### ChIP-Rx analysis

Reads were aligned to the human genome hg38 and mouse reference genome mm10 using Bowtie v2.10^103^. A normalization factor was generated by establishing the number of human reads relative to spike-in mouse genome. Ambiguous reads aligning to more than one reference genome and duplicated reads were removed. Bigwig files were generated at 10bp resolution using the bamCoverage utility from the deepTools2 suite^108^ and the data was visualized using the UCSC genome browser. Peaks were called using MACS2^104^ with FDR <0.05 and annotated using HOMER^110^.

#### RNA-seq analysis

For each pair of FASTQ files, BBDuk from BBTools^109^ package was used to trim adaptors and low-quality reads. Filtered reads were aligned to the hg38 homo sapiens reference genome (GRCh38.98) using HISAT2^112^ (version 2.2.1). The SAMtools program^105^ was used to convert SAM files to sorted BAM files. Next, featureCounts^113^ (version 1.6.4) was used to count reads per gene, which had been assigned to gene identifier features annotated by the Ensembl 98 hg38 genome annotation. The resulting read counts table was imported into R, where the DESeq2 package^106^ (version 1.48.0) was used to identify differentially expressed genes with a fold-change of >1.5, a baseMean expression of >20, and adjusted p values <0.05. Adjusted p values were calculated in DESeq2 using the Benjamini–Hochberg (BH) procedure to control the false discovery rate (FDR). Differential expression results were visualized using volcano plots. Genes were highlighted in red if they met all of the following criteria: adjusted p-value < 0.05, minimum 1.5-fold change, and base mean expression > 20. Non-significant genes are shown in grey. Plots were generated using ggplot2^111^.

#### QuantSeq analysis

Raw sequencing reads were aligned to the hg38 homo sapiens reference genome (GRCh38.98) using HISAT2^112^ and the aligned files were converted to bigwig files using bamcoverage^117^. Sequence reads were aggregated into a count for each gene using featurecounts^113^ and differentially expressed genes were identified using DESeq2^106^. Volcano plots of differentially expressed genes were generated using EnhancedVolcano^114^.

## ACKNOWLEDGMENTS

We thank members of the Bracken lab for helpful discussions throughout the development of this work. We would also like to thank Evan Healy, Orla Deevy, Jingjing Li, and Karsten Hokamp for their help in preparing this manuscript. Work in the Bracken lab was supported by the Worldwide Cancer Research and Irish Cancer Society charities (24-0350), an Irish Research Council Advanced Laureate Award (IRCLA/2019/21) and SFI, under the SFI Investigators program (SFI/16/IA/4562). Work in the Conway lab was supported by a Wellcome Trust Early Career Award [225152/Z/22/Z], a Worldwide Cancer Research award [225152/Z/22/Z] and a Research Ireland Pathway Program grant (21/PATH-S/9384). UCD mass spectrometry was supported by The Comprehensive Molecular Analytical Platform (CMAP) under The Science Foundation Ireland (SFI) Research Infrastructure Programme, reference 18/RI/5702. D.A. was supported by a PhD fellowship from the Irish Cancer Society Translational Research Scholarship (CRS21ANG). J.N. was supported by a PhD fellowship from the Irish Research Council Government of Ireland Postgraduate Scholarship Programme (GOIPG/2019/2784). G.H. was supported by a PhD fellowship from the Irish Research Council Government of Ireland Postgraduate Scholarship Programme (GOIPG/2023/4606). D.N. was supported by a PhD fellowship from Research Ireland through the Research Ireland Centre for Research Training in Genomics Data Science under Grant number 18/CRT/6214.

## AUTHOR CONTRIBUTIONS

D.A., A.P.B., E.C., and J.N. conceived the study. D.A. performed the genome-wide CRISPR screens and validations, PRC2 purifications, all sgRNA-based epigenomic and transcriptomic experiments, and some of the bioinformatical analyses. J.N. and E.C. performed the PRC2-tiling screen in lymphoma cells. J.N. performed all validations and all shRNA-based and pharmacological epigenomic and transcriptomic experiments. M.M. performed the PRC2-tiling screen in MRT cells. E.D. performed mass spectrometry analyses. A.F. provided structural guidance to identify zinc-finger residues for targeted mutation. D.N., C.M. and C.W. contributed to computational analyses of proteomic and genomic datasets. J.N., E.L. and A.M. performed the bioinformatical analysis of the PRC2-tiling screens. E.C., G.L.B., and C-W.C. contributed to the design of the PRC2-tiling CRISPR screening experiments. D.R., G.H., and M.D. supported bench-based experimental work. D.A. and A.P.B wrote the first draft of the manuscript, and all authors contributed to its subsequent writing and critical review.

---

**Figure S1.**
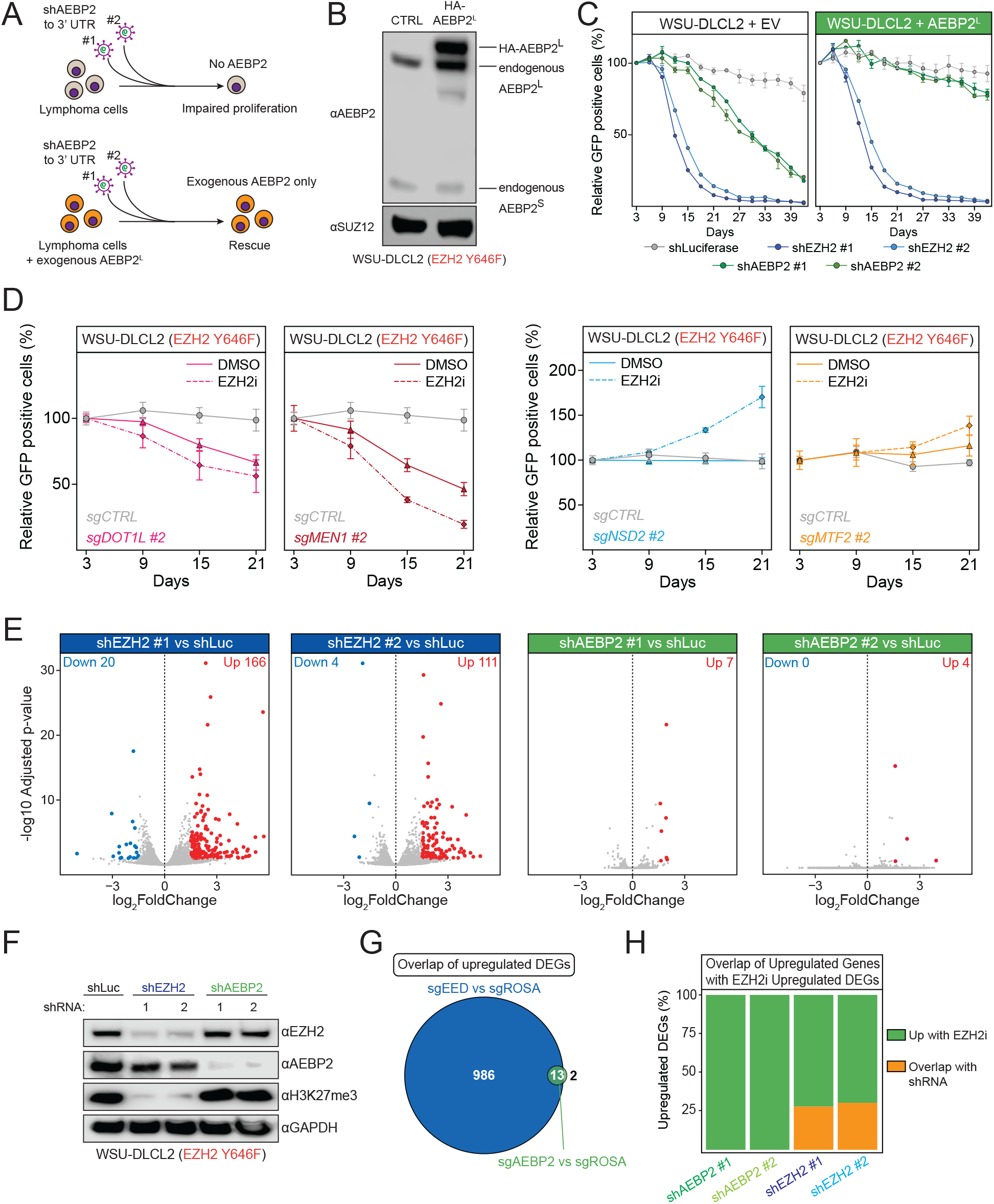
AEBP2 is an EZH2 co-dependency in B-cell lymphoma but dispensable for global H3K27me3 and gene repression. **A**. Schematic illustrating the shRNA-based rescue strategy. Two independent shRNAs targeting the 3’ UTR of AEBP2, effectively knock down endogenous AEBP2, while leaving exogenous AEBP2^L^, lacking the 3′ UTR, unaffected. Immunoblot of HA-tagged exogenous AEBP2 expression in WSU-DLCL2 (EZH2-mutant) lymphoma cells. **B**. Growth competition assays in WSU-DLCL2 (EZH2-mutant lymphoma) cells expressing wild-type AEBP2^L^ compared to empty vector, transduced with shRNAs targeting the indicated genes (n=3, data represents mean ± SD). **C**. Growth competition assays in Cas9-expressing WSU-DLCL2 (EZH2-mutant lymphoma) cells transduced with sgRNAs targeting the indicated genes followed by treatment with low dose EZH2 inhibitor (Tazemetostat) (n=3, data represents mean ± SD). **D**. Volcano plots depicting differentially expressed genes in WSU-DLCL2 (EZH2-mutant lymphoma) cells following shRNA-mediated knock-down of EZH2 or AEBP2. The number of genes differentially expressed both up (red) and down (blue) are indicated on the plots (n=3 biological replicates; significance cutoff: p < 0.05; minimum 1.5-fold change). **E**. Immunoblots of the indicated proteins and histone modifications in WSU-DLCL2 (EZH2-mutant lymphoma) cells transduced with two independent EZH2- and AEBP2-targeting shRNAs. **F**. Venn diagram demonstrating overlap of differentially expressed genes upregulated (DEGs up) in WSU-DLCL2 lymphoma cells with CRISPR Cas9-mediated KO of *EED* or *AEBP2* compared to control (*ROSA*). **G**. Bar plots depicting degree of overlap (orange) between upregulated genes in WSU-DLCL2 (EZH2-mutant lymphoma) cells treated with EZH2i compared to EZH2 or AEBP2 knockdown.

**Figure S2.**
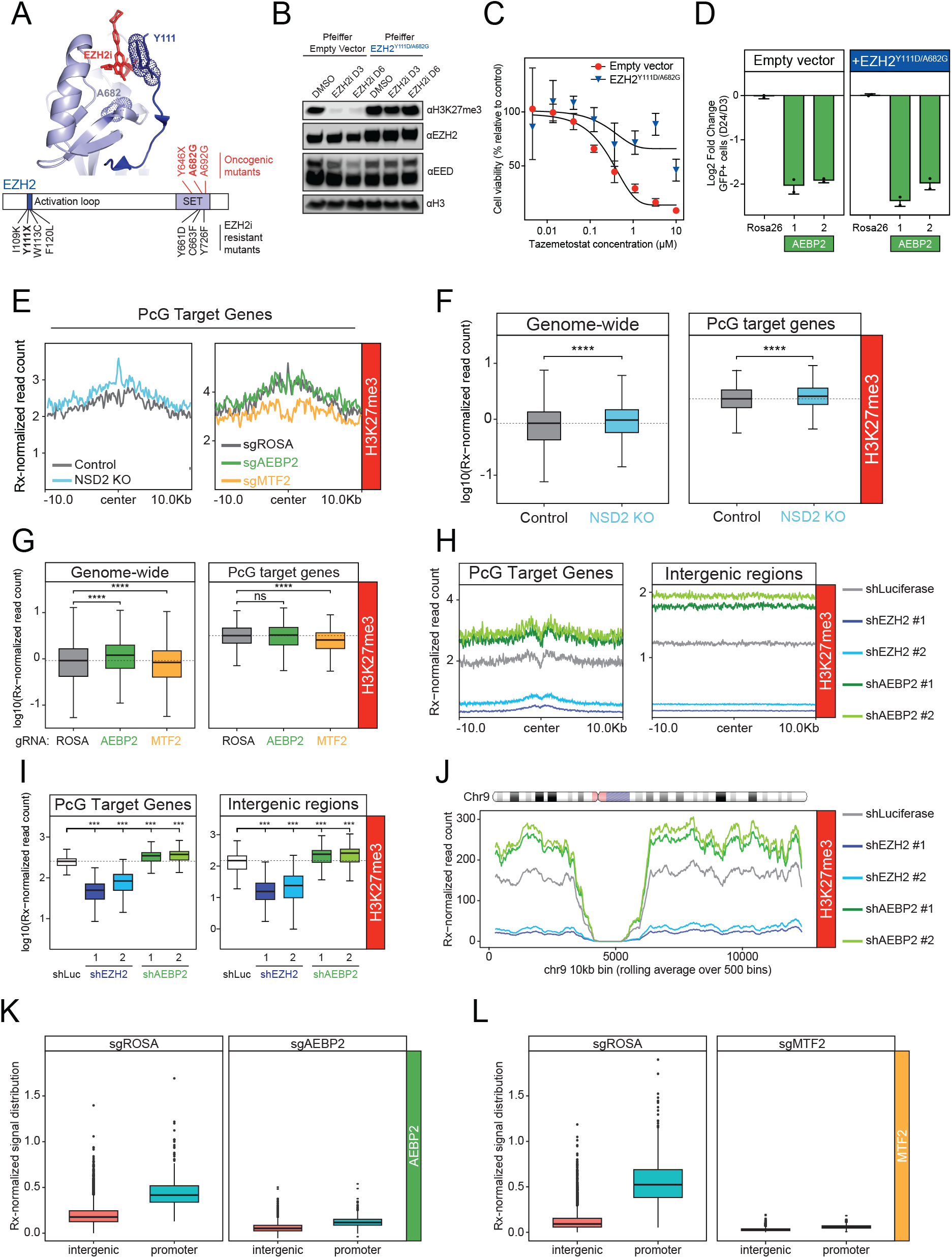
AEBP2 and NSD2 loss both increase H3K27me3 despite conferring opposing sensitivity to EZH2 inhibitors. **A**. Modelling of PRC2 enzymatic inhibitor drug binding to EZH2 in its active conformation, with EZH2 inhibitor binding residue Y111X (dots) and EZH2 oncogeneic mutant A682G annotated^94^ (Top). Schematic representation of EZH2 with hotspot oncogenic mutations and mutations conferring drug resistance in lymphoma cell lines highlighted (Bottom). **B**. Immunoblots of the indicated PRC2 proteins and histone modifications in the EZH2-mutant cell line Pfeiffer expressing empty vector or EZH2^Y111D/A682G^ treated with either DMSO or EZH2 inhibitor Tazemetostat (30nM) for 3 or 6 days. **C**. Cell viability assay using the CellTiter-Glo® reagent in Pfeiffer (EZH2-mutant lymphoma) empty vector (EV) or EZH2^Y111D/A682G^ cell lines treated with indicated doses of Tazemetostat for 6 days. **D**. Growth competition assays in Cas9-expressing Pfeiffer (EZH2-mutant lymphoma) empty vector (EV) or EZH2Y^111D/A682G^ cell lines transduced with sgRNAs targeting the indicated genes (n=3, data represents mean ± SD). **E**. Average plots showing H3K27me3 CUT&RUN-Rx enrichment at genomic windows +/-10kb from PRC2-bound promoters in control and *NSD2* KO WSU-DLCL2 (EZH2-mutant lymphoma) cells (left) and in Cas9-expressing WSU-DLCL2 (EZH2-mutant lymphoma) cells transduced with sgRNAs targeting the indicated PRC2 genes (right). TES and TSS indicate Transcription End Site and Transcription Start Site respectively. **F**. Boxplots of H3K27me3 CUT&RUN-Rx signal at genome-wide 10kb bins and at PRC2-bound promoters in control and *NSD2* KO WSU-DLCL2 (EZH2-mutant lymphoma) cells. Significance determined using pairwise Wilcoxon rank-sum tests, ****p value ≤ 0.0001. **G**. Boxplots of H3K27me3 CUT&RUN-Rx signal at genome-wide 10kb bins and at PRC2-bound promoters in Cas9-expressing WSU-DLCL2 (EZH2-mutant lymphoma) cells transduced with sgRNAs targeting the indicated PRC2 genes. Significance determined using pairwise Wilcoxon rank-sum tests, ****p value ≤ 0.0001. **H**. Average plots showing H3K27me3 ChIP-Rx enrichment at genomic windows +/-10kb from PRC2-bound promoters (left) and at all intergenic sites (right) in WSU-DLCL2 (EZH2-mutant lymphoma) cells transduced with two independent EZH2- and AEBP2-targeting shRNAs. **I**. Boxplots of H3K27me3 ChIP-Rx signal at PRC2-bound promoters and intergenic regions in WSU-DLCL2 (EZH2-mutant lymphoma) cells transduced with two independent EZH2- and AEBP2-targeting shRNAs. Significance determined using pairwise Wilcoxon rank-sum tests, ***p value ≤ 0.001. **J**. Rolling average plots presenting H3K27me3 ChIP-Rx signal across the whole of chromosome 9 in WSU-DLCL2 (EZH2-mutant lymphoma) cells with two independent EZH2- and AEBP2-targeting shRNAs. Visualized using 10kb genomic windows. **K**. Boxplots comparing the distribution of AEBP2 CUT&RUN-Rx signal at PRC2-bound promoters and intergenic regions in Cas9-expressing WSU-DLCL2 (EZH2-mutant lymphoma) cells transduced with sgRNAs targeting the indicated genes. **L**. Boxplots comparing the distribution of MTF2 CUT&RUN-Rx signal at PRC2-bound promoters and intergenic regions in Cas9-expressing WSU-DLCL2 (EZH2-mutant lymphoma) cells transduced with sgRNAs targeting the indicated genes.

**Figure S3.**
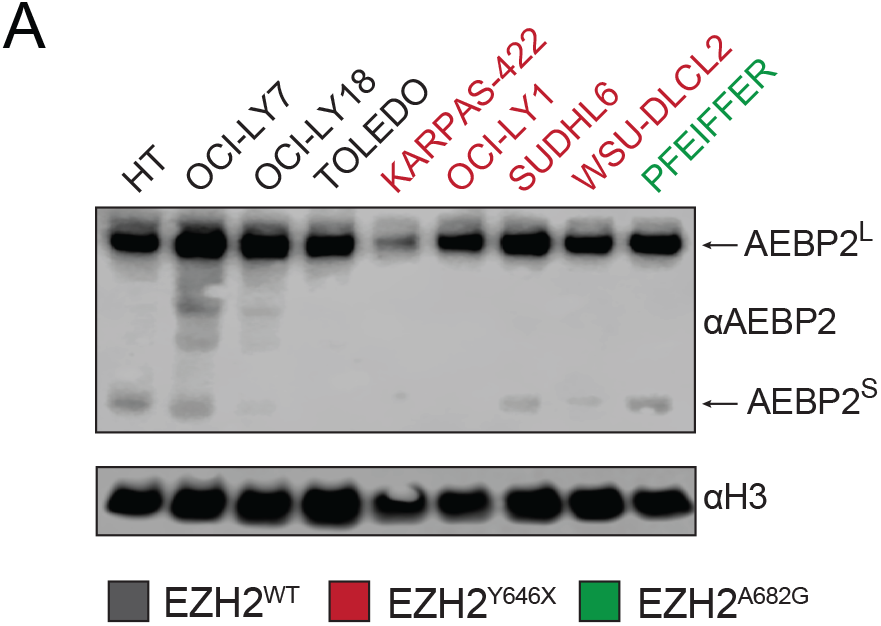
AEBP2 confers its essential function in EZH2-mutant lymphoma in a PRC2 complex lacking JARID2. **A**. Immunoblots of AEBP2 and histone H3 in a panel of germinal center B-cell lymphoma cell lines with the EZH2 genotypes color coded as grey (wild-type), red (Y646X), and green (A682G).

**Figure S4.**
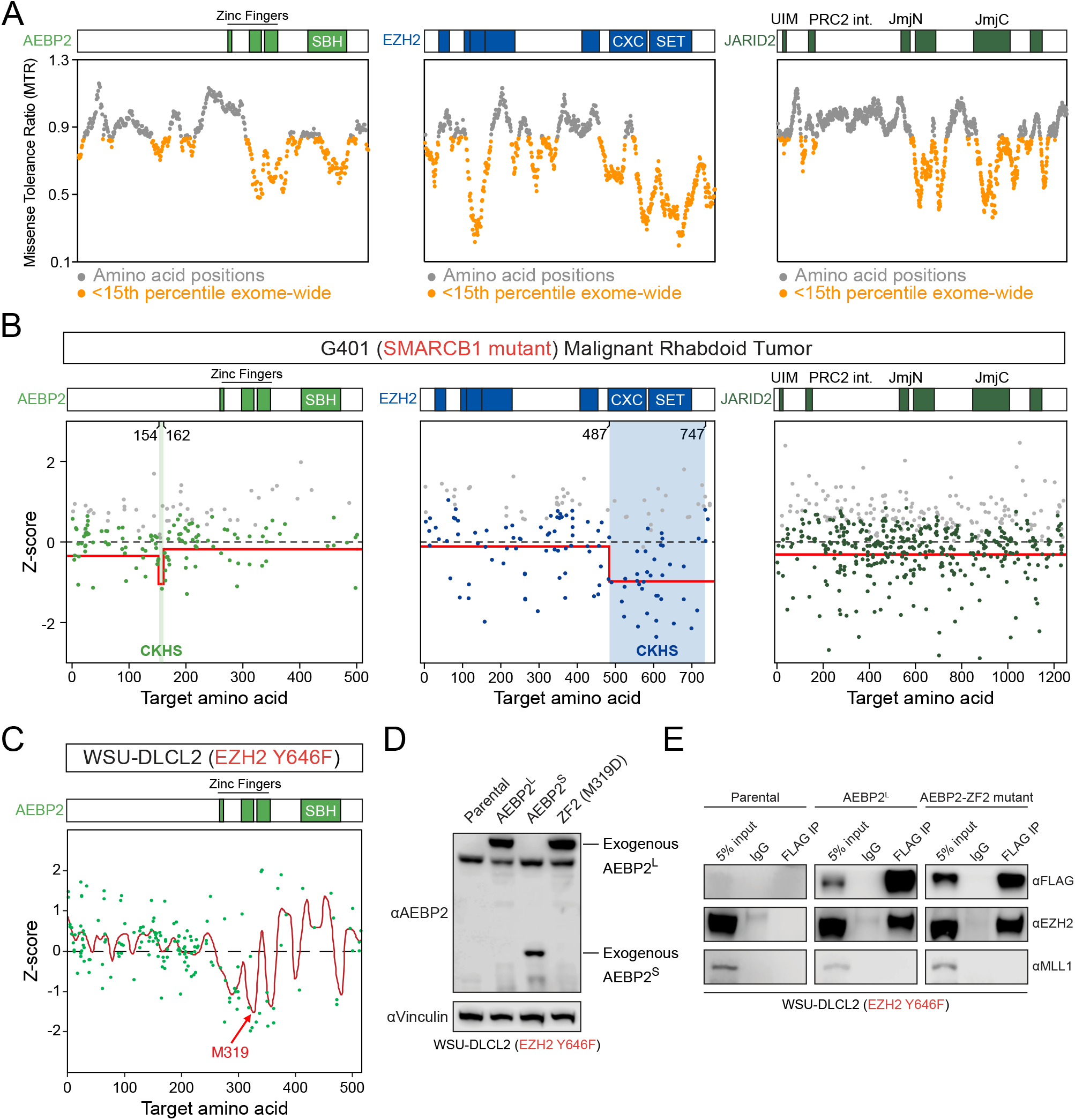
AEBP2 chromatin binding via zinc fingers is critical for its role in lymphoma. **A**. Missense Tolerance Ratio plots representing regional intolerance to missense mutations using the RGC-ME database of missense mutations in healthy individuals (https://rgc-research.regeneron.com/me/home)^95^. We used a sliding window of 31 amino acids across AEBP2, EZH2 and JARID2. Amino acids in the top 15% of those intolerant to missense mutations genome wide are colored orange. The relevant domains of each protein are indicated. SBH indicates the SUZ12-binding helix domain. **B**. CRISPR PRC2-tiling screen results of AEBP2, EZH2 and JARID2 in G401 (SMARCB1-mutant MRT) cells (n=2 biological replicates). sgRNAs are indicated by dots mapped to the target amino acid. CRISPR-knockout hyper-sensitivity (CKHS) regions are indicated in the shaded regions. The average depletion of sgRNAs targeting a protein segment is indicated by the red line (Z-score). The relevant domains of each protein are indicated. SBH indicates the SUZ12-binding helix domain. **C**. CRISPR-tiling screen results of AEBP2 in WSU-DLCL2 (EZH2-mutant lymphoma) cells (n=2 biological replicates) with sgRNAs indicated by dots mapped to the target amino acid and analysis carried out using the “smoothen” model^96^. The average depletion of sgRNAs targeting a protein segment is indicated by the red line (Z-score). The relevant domains of each protein are indicated. SBH indicates the SUZ12-binding helix domain. The zinc finger 2 M319 residue is highlighted. **D**. Immunoblots of the indicated proteins in WSU-DLCL2 (EZH2-mutant lymphoma) cells expressing the HA/FLAG-tagged AEBP2 wild-type and mutant constructs. **E**. Immunoblots of the indicated proteins in FLAG protein co-immunoprecipitations in WSU-DLCL2 (EZH2-mutant lymphoma) cells expressing the HA/FLAG-tagged AEBP2 wild-type and mutant constructs.

**Figure S5.**
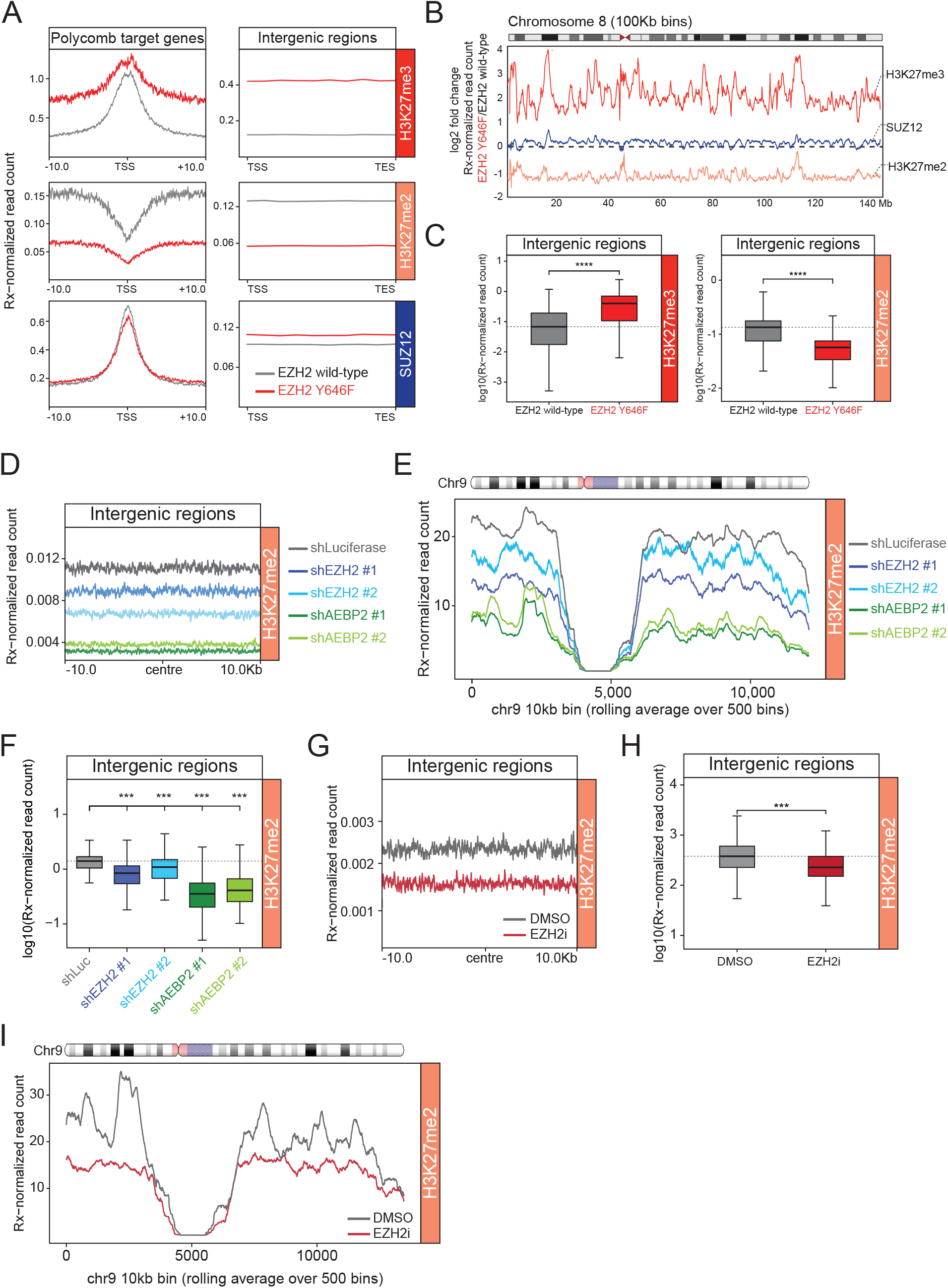
AEBP2 sustains residual H3K27me2 essential for continued proliferation of EZH2-mutant lymphoma. **A**. Average plots showing H3K27me2, H3K27me3, and SUZ12 ChIP-Rx enrichment at all intergenic sites in OCI-LY7 (EZH2 wild-type lymphoma) cells expressing either wild-type EZH2 or the mutant EZH2 Y646F. **B**. Rolling average plots presenting Log_2_ Fold-Change of ChIP-Rx signal for H3K27me2, H3K27me3, and SUZ12 across the whole of chromosome 9 between OCI-LY7 (EZH2 wild-type lymphoma) cells expressing either wild-type EZH2 or the mutant EZH2 Y646F. Visualized using 100kb genomic windows. **C**. Boxplots of H3K27me2 and H3K27me3 ChIP-Rx signal at all intergenic regions in OCI-LY7 (EZH2 wild-type lymphoma) cells expressing either wild-type EZH2 or the mutant EZH2 Y646F. Significance determined using pairwise Wilcoxon rank-sum tests, ****p value ≤ 0.0001. **D**. Average plots showing H3K27me2 ChIP-Rx enrichment at all intergenic sites in WSU-DLCL2 (EZH2-mutant lymphoma) cells transduced with two independent EZH2- and AEBP2-targeting shRNAs. **E**. Rolling average plots presenting H3K27me2 ChIP-Rx signal across the whole of chromosome 9 in WSU-DLCL2 (EZH2-mutant lymphoma) cells transduced with two independent EZH2- and AEBP2-targeting shRNAs. Visualized using 10kb genomic windows. **F**. Boxplots of H3K27me2 ChIP-Rx signal at all intergenic regions in WSU-DLCL2 (EZH2-mutant lymphoma) cells transduced with two independent EZH2- and AEBP2-targeting shRNAs. Significance determined using pairwise Wilcoxon rank-sum tests, ***p value ≤ 0.001. **G**. Average plots showing H3K27me2 ChIP-Rx enrichment at all intergenic sites in WSU-DLCL2 (EZH2-mutant lymphoma) cells treated with EZH2 inhibitor (Tazemetostat) compared to control (DMSO). **H**. Boxplots of H3K27me2 ChIP-Rx signal at all intergenic regions in WSU-DLCL2 (EZH2-mutant lymphoma) cells treated with EZH2 inhibitor (Tazemetostat) compared to control (DMSO). Significance determined using pairwise Wilcoxon rank-sum tests, ***p value ≤ 0.001. **I**. Rolling average plots presenting H3K27me2 ChIP-Rx signal across the whole of chromosome 9 in WSU-DLCL2 (EZH2-mutant lymphoma) cells treated with EZH2 inhibitor (Tazemetostat) compared to control (DMSO). Visualized using 10kb genomic windows.

## KEY RESOURCES TABLE

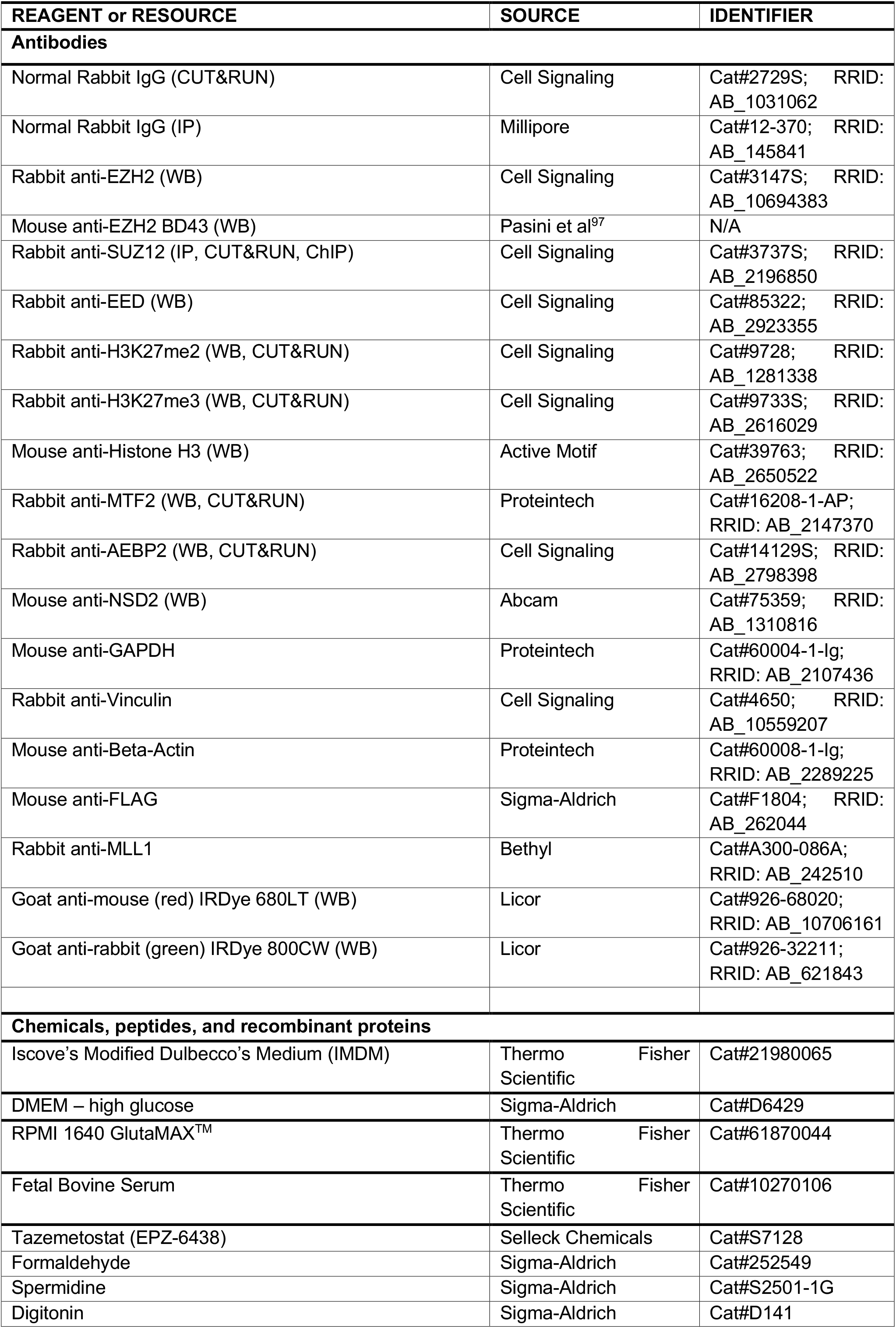
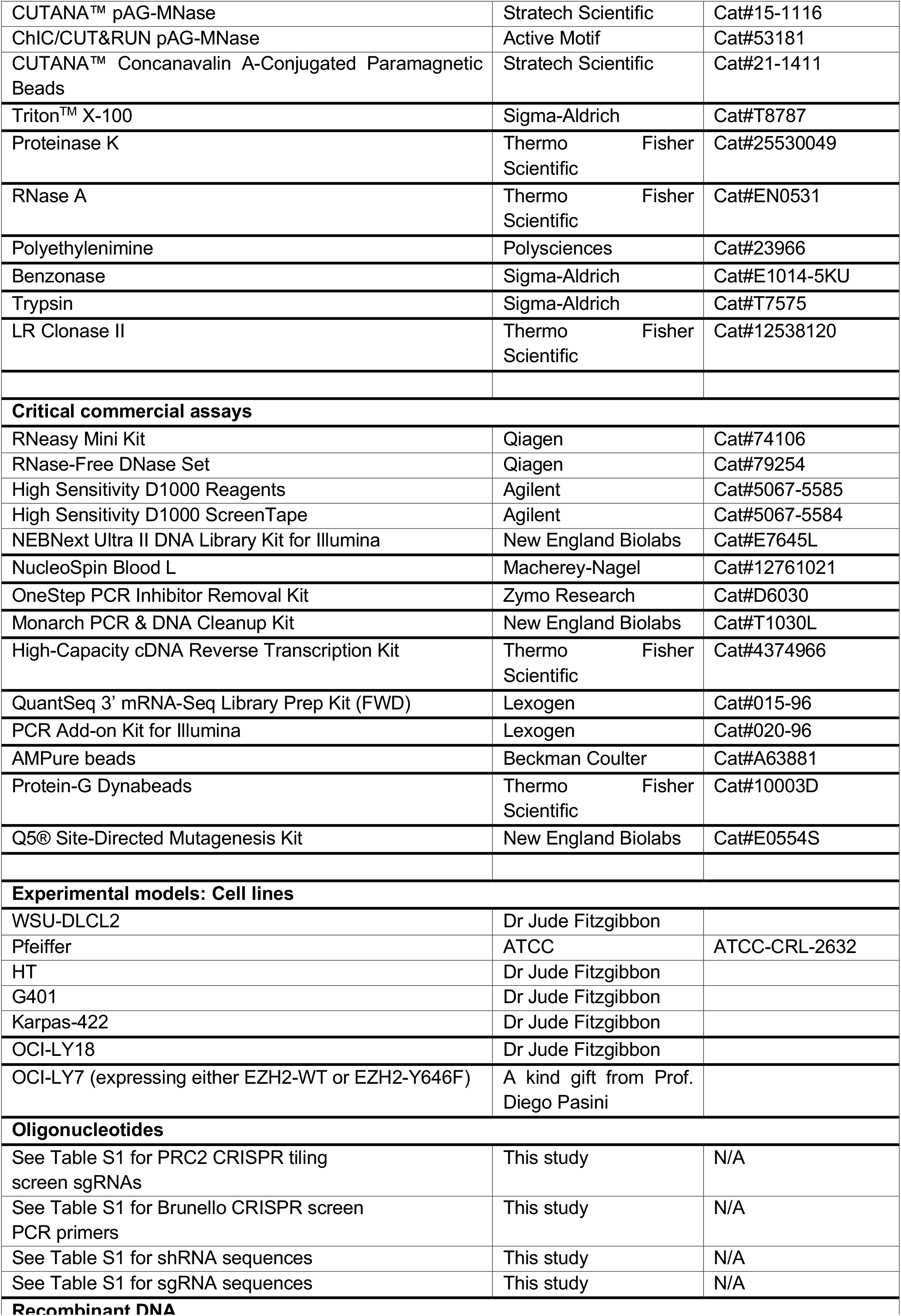
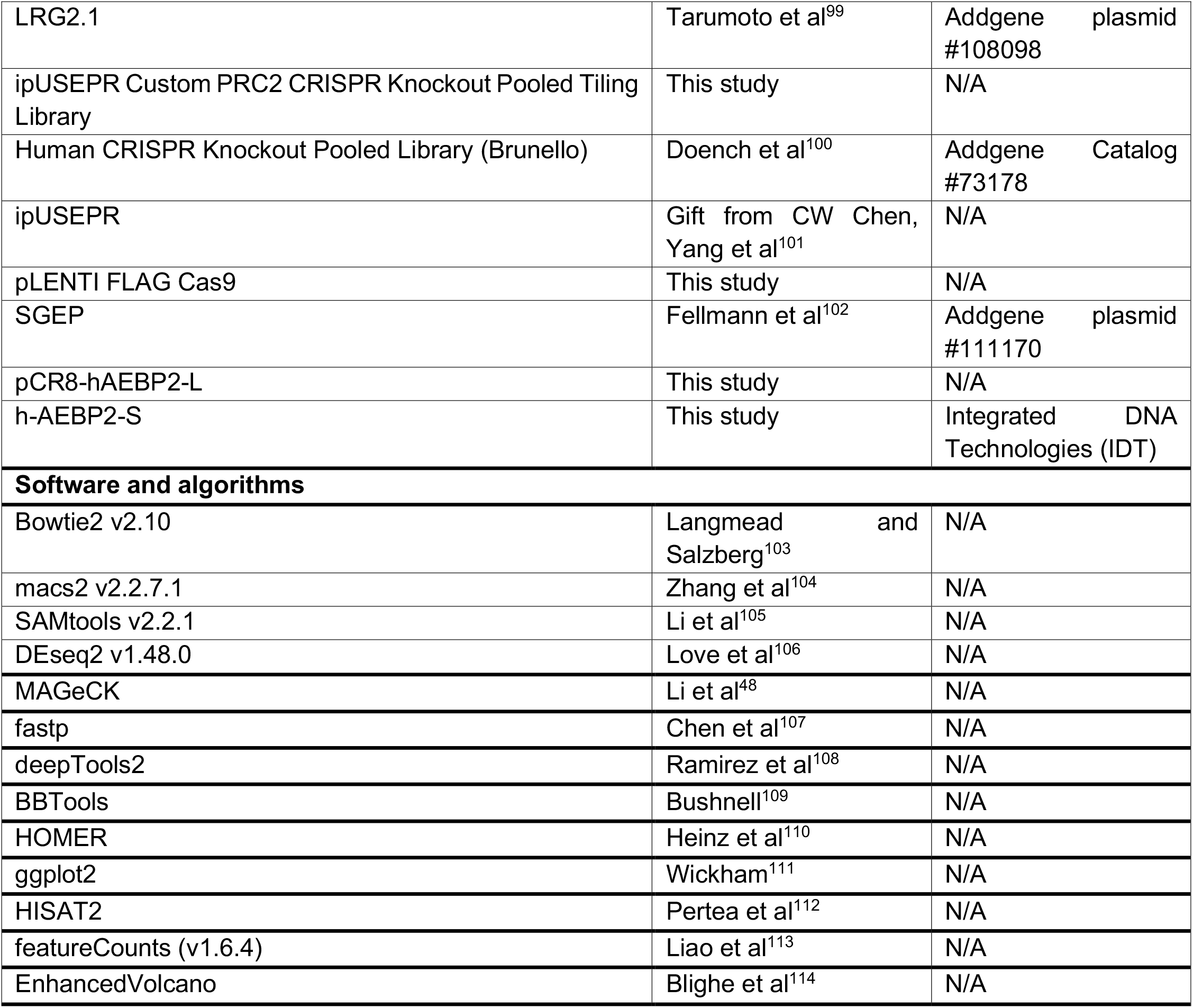

